# Cross-Species Efficacy of Combinatorial Gene Therapy for Osteoarthritis and Correction of Neuro-Inflammatory Pain Mechanisms

**DOI:** 10.64898/2026.07.27.740924

**Authors:** Erica Secor, Jun Wang, Zelong Dou, Jiansen Yan, Camilla Majano, Matthew Woodman, Oscar Ruiz, Joud Al Azaat, Duncan Crosby, Racel Cela, Sarah Pownder, Julie B Engiles, Donna Palmer, Mingming Jiang, Carolina Leynes, Ismail Yaman, Mira Jeong, Gerhard Sponder, Stanislav Plutziki, Lisa Yuva, Surabi Veeraragavan, Russell S. Ray, Joshua D. Wythe, Benjamin R. Arenkiel, Rui Chen, Kim C. Worley, RE-JOIN Consortium, Philip Ng, Masataka Suzuki, Kilian Guse, Yangjin Bae, Nele A. Haelterman, Heidi Reesink, Brendan Lee

## Abstract

Osteoarthritis is a leading cause of chronic pain and disability, which lacks disease-modifying treatment. Given the complex multi-tissue and multifactorial drivers behind disease progression, effective treatments will require simultaneously targeting several mechanisms underlying joint degeneration and pain. Here, we developed and evaluated a combinatorial gene therapy, consisting of a high-capacity adenoviral vector carrying two therapeutic genes to target distinct pathological mechanisms: inflammation (IL-1Ra) and chondrocyte health (PRG4). Intra-articular delivery of this treatment improved functional, structural, and pain outcomes in murine and equine osteoarthritis models. In addition, treatment normalized inflammatory environments in joint tissues, as well as in the dorsal root ganglia (DRG) known to harbor joint-innervating sensory neurons. Moreover, gene therapy reversed OA-induced molecular signatures of neural hyperexcitability, suggesting amelioration of peripheral sensitization. Collectively, these findings support combinatorial gene therapy as a promising treatment for osteoarthritis, while identifying neuroinflammatory signatures for correction of disease progression and pain.

**One Sentence Summary:** A single intra-articular injection of a combinatorial gene therapy slows OA progression and reduces pain in small and large animal models.

## INTRODUCTION

Osteoarthritis (OA) is the most common form of arthritis in adults and a leading cause of disability and chronic pain worldwide, for which no disease-modifying treatment currently exists (*1–3*). Accordingly, disease management is centered around symptomatic relief to improve chronic pain and joint function, with surgical interventions reserved as a last resort for patients with treatment-resistant symptoms that affect quality of life (*4*). Initially considered a wear-and-tear disease marked primarily by articular cartilage loss, OA is now understood to be a complex, multifactorial condition that affects the whole joint organ, including its immunological and neurological components such as the innervating dorsal root ganglia (DRG)(*2, 5*). Hence, it is becoming increasingly clear that effective OA therapeutic approaches for OA must simultaneously target multiple drivers of disease, an emerging theme for complex diseases.

Numerous preclinical models have been developed to study pathogenic mechanisms, identify drivers of disease progression and pain, and screen potential therapeutic candidates (*6*). Such approaches primarily leverage rodent animal models, spanning genetic models that spontaneously develop OA, post-traumatic OA (PTOA) models that use surgical (e.g. destabilization of the medial meniscotibial ligament (DMM) or anterior cruciate ligament transection (ACLT) murine models) or non-invasive methods to decrease joint stability, or inflammatory models that chemically induce joint degeneration (*6–10*). While rodent models have yielded key insights into OA disease mechanisms, several factors, including biomechanical and anatomical differences with humans, may limit the translational potential of rodent-focused research studies (*7*). Given OA’s complex nature, studies that employ multiple animal models, with orthogonal validation across different species, will likely yield a deeper, more impactful and translatable understanding of underlying pathogenic mechanisms that can be targeted for meaningful disease modification in patients. Hence, there is a demonstrable need for studies in large animal models to bridge findings between tractable rodent models and humans. Equine models are particularly well suited to fill this role, as their joint anatomy, biomechanics, and articular cartilage are the most comparable to humans, and horses are predisposed to naturally develop OA (*9, 11*). The carpal osteochondral fragment model is the most frequently cited equine surgical OA model, inducing progressive OA in the absence of joint instability, unlike other preclinical surgical OA modalities (*11–14*). This approach consistently induces hallmark features of OA pathogenesis, including progressive cartilage matrix breakdown, synovial inflammation, and biochemical changes in synovial fluid, and has been widely used to evaluate therapies with strong translational relevance to human OA (*12, 15, 16*). However, whether the neuroinflammatory signatures that have emerged as critical drivers of OA in murine models and humans are also present in equine OA is unknown.

An extensive body of work has identified low-grade, chronic inflammation as an important driver of OA progression (*17–20*). Pursuant to these findings, studies in small and large animal models of OA showed that dampening inflammation by inhibiting interleukin 1 (IL-1) signaling effectively reduces pain and disease progression, which informed multiple clinical studies (*14, 16, 21–23*). Importantly, mixed results from preclinical and clinical studies with various IL-1 modulators to date suggest that sustained inhibition of this pathway may be detrimental to joint health (*21, 24–26*). Hence, developing therapeutic strategies that enable regulated expression of signaling modifiers may be important for complex disorders, such as osteoarthritis.

High-Capacity Adenovirus (HCAd)-based gene therapy holds great promise for the treatment of chronic joint diseases. Its local, intra-articular administration confers low immunogenicity while its high transduction efficiency yields stable, long-term expression of therapeutic targets from cells within the synovium, cruciate ligaments, and cartilage (*27, 28*). Unlike AAV, HCAd’s large genetic payload capacity (∼4.2kb vs ∼36kb) allows combining multiple therapeutic transgenes into a single vector. This property is particularly relevant if use and combination of large genes is desired. We previously developed an HCAd-based gene therapeutic approach to conditionally express the IL-1 receptor antagonist (IL-1Ra) in the presence of inflammation through regulated expression with an NFκB promoter (*HCAd-NFκB::IL1-Ra*)(*14*). This approach effectively slows disease progression in murine and equine animal models of OA, and a recently concluded phase I clinical study showed sustained pain relief up to three years following a single injection (NCT06884865) (*29–31*). These results demonstrate that intra-articular, inflammation-regulated IL-1Ra expression reduces pain and disease progression while improving motor function.

Parallel research identified additional mechanisms underlying OA pathogenesis, including chondrocyte homeostasis, matrix degradation, and others (*5, 32*). These studies identified additional promising therapeutic targets, such as lubricin (Proteoglycan 4; *Prg4*), a lubricating chondroprotective and joint homeostatic glycoprotein whose genetic loss causes early onset OA in mice and humans (*33, 34*). We previously showed that HCAd-mediated Prg4 expression (*HCAd-EF1::Prg4*) slows OA disease progression in multiple murine OA models (*28, 35*). Moreover, co-injection of distinct *IL-1Ra* and *Prg4* vectors into murine OA joints delays disease progression more compared to injection of either vector alone (*35*), demonstrating an added benefit of targeting multiple disease mechanisms for OA.

Here, we generated a single HCAd gene therapy that delivers both therapeutic genes (IL-1Ra and PRG4; i.e., dual vector) and show its effective modification of pain and structural phenotypes in rodent and equine OA models. We assessed the dual vector’s ability to modulate OA disease pathogenesis using a battery of histological, molecular, and single nucleus transcriptomic assays. Critically, surgically induced OA alters the neuro-immune landscape in murine and equine synovial tissues and in the joint-innervating equine dorsal root ganglia (DRG), and dual vector treatment reversed these changes. The behavioral, pain, and structural improvements across the models correlated with normalization of the immune and neuronal bio-signature in joint tissues and DRGs. In sum, our studies suggest that a HCAd-mediated expression of both PRG4 and inflammation-induced IL-1Ra is a promising therapeutic combination for clinical translation. Moreover, these studies identify an immune and neuronal signature that is associated with OA that can be normalized by this clinically relevant treatment approach.

## RESULTS

### Intra-articular dual vector treatment effectively slows joint disease progression in murine OA models

Given the robust treatment effects of HCAd-based gene therapeutic strategies in murine and equine models of OA (*14, 28, 35*), the promising results of HCAd-based IL-1Ra gene therapy in a recently concluded phase I trial (NCT06884865)(*29, 30*), and our previous study showing further improved protection when co-injecting HCAd-vectors separately expressing IL-1Ra and Prg4 (*35*), we leveraged HCAd’s large genetic payload capacity to combine two promising therapeutic genes (*IL-1Ra* and *Prg4*) in the same vector (dual vector: *HCAd-NFkb::IL-1Ra; EF1::Prg4*; **Suppl Fig S1a,c,e**). We first tested if this dual vector could similarly enhance therapeutic efficacy compared to injection of each vector alone in a rat ACLT model. 9- to 10-week-old Sprague Dawley rats were subjected to ACLT surgery and were injected with vector formulation (vehicle), *HCAd-NFkb:rIL-1Ra*, *HCAd-EF1:rPRG4*, or with *HCAd-NFkb::rIL-1Ra; EF1::rPrg4* (rat dual vector) 1 week post-surgery (**Suppl Fig S1b)**. Histopathological scoring of knee joints, collected 15 weeks post-surgery validated our previous findings, showing enhanced protection against ACLT-induced joint degeneration upon combining both therapeutic genes in the lateral compartment based on standard histopathological scoring (**Suppl Fig S2**).

We next expanded our studies to a second OA model (mouse DMM model) to study the dual vector’s effects on inflammation, cartilage degeneration, and pain, in greater detail, and to explore potential underlying mechanisms. *In vitro* validation of the murine dual vector in 293T cells showed comparable genomic copy numbers among different HCAd vectors, 2 days after transduction (**Suppl Fig S3a**). TNFα-stimulation of transduced cells induced IL-1Ra mRNA and protein expression compared to non-stimulated cells, confirming inflammation-dependent transgene expression from the *NFkB* promoter in both the mono-vector (*HCAd-NFkb::IL-1Ra*; **Suppl Fig S3b,d**) and the dual vector (**Suppl Fig 3c,e**). In parallel, PRG4 transcript levels were significantly increased compared to non-transduced cells, reflecting constitutive EF1-promoter activity (**Suppl Fig S3j**).

We next tested the efficacy of intra-articular injection of the dual vector for delaying OA progression in the murine DMM surgical OA model, which induces pain behaviors and mild-to-moderate OA at 3 months post-surgery (*6, 8, 9*). DMM mice, which received intra-articular (IA) injection of murine dual vector (1×10^8 vp/ knee) or PBS (vehicle control) two weeks post-surgery, and sham mice (surgical control where the joint capsule is opened and closed without inducing joint damage) were collected 12 weeks post-surgery (**Suppl Fig S1d, S4a**). OA pathology was assessed by phase-contrast microcomputed tomography (µCT, **Fig 1a,b**) and standard histopathological scoring (OARSI; **Suppl Fig S4b,c**). Femoral articular cartilage volumes were significantly greater in dual vector-treated OA joints compared to vehicle-treated OA joints, and articular cartilage volumes of sham control joints were not significantly different from dual vector-treated joints (**Fig 1a,b**). Consistent with the volumetric change of articular cartilage, OARSI scores in the dual vector group trended lower compared to vehicle-treated controls (**Suppl Fig S4b,c**). Collectively, these morphological data indicate that dual vector treatment effectively attenuates injury-induced articular cartilage loss in DMM mice.

**Fig. 1.**
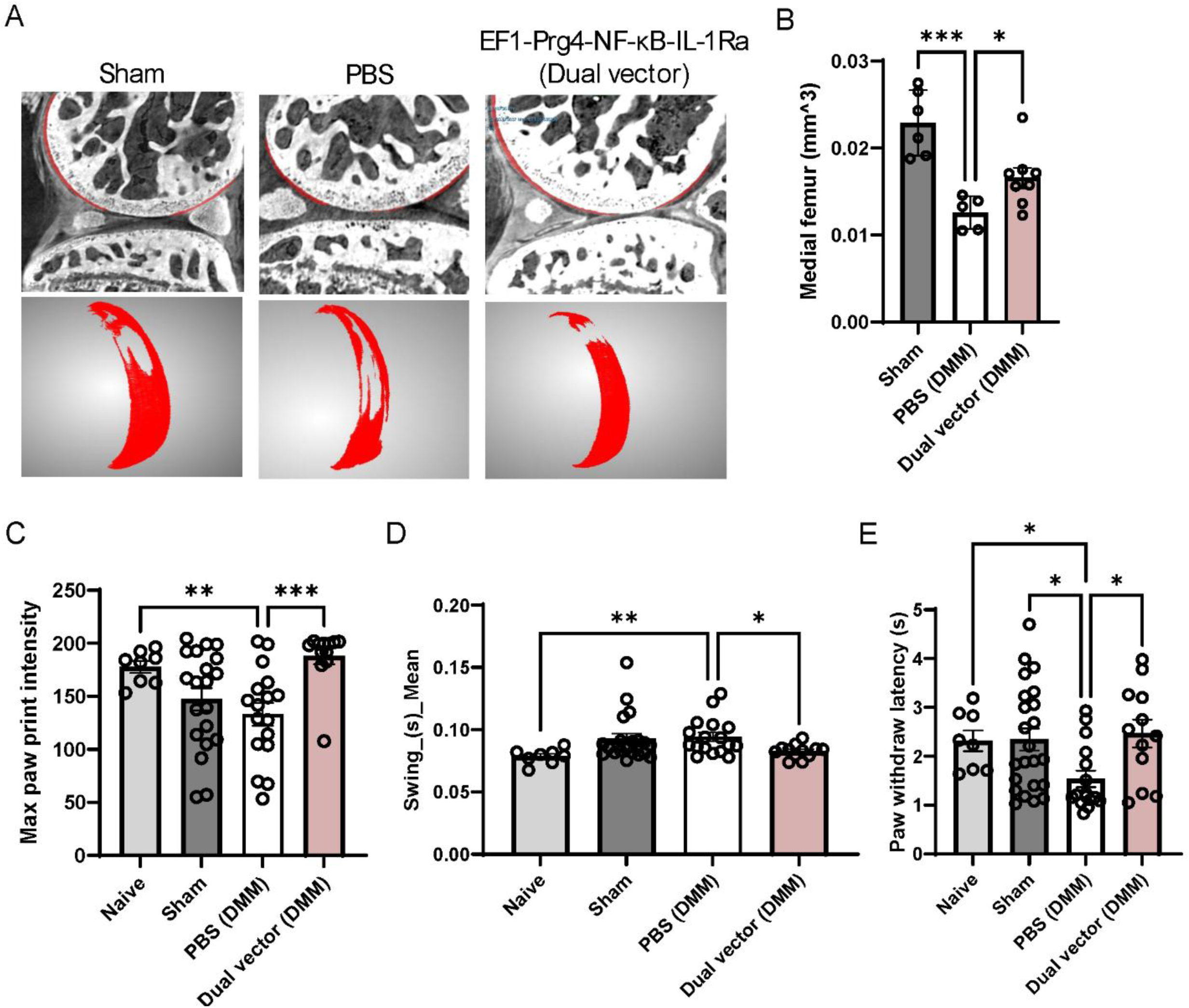
Efficacy of dual vector in murine OA. Results of the murine surgical OA model treated with *HCAd-NFkb::mIL-1Ra; EF1::mPrg4* (dual vector) or vehicle control (PBS). (A) Representative sagittal phase-contrast µCT image of the mouse knee joint (top), frontal view of digitally segmented medial femoral articular cartilage shown in red (bottom). (B) Quantification of segmented medial femoral articular cartilage volume. (C) CatWalk-based quantification of paw print intensity. (D) CatWalk-based quantification of swing time. (E) Hot plate assay measuring mild response latency to thermal stimulation. Statistical significance was assessed using Brown–Forsythe and Welch ANOVA with Dunnett’s T3 multiple-comparisons test. *: p<0.05, **: p<0.01, *** = p<0.001. Mean values ±SEM are shown.

OA progression is associated with inflammation, pathological whole joint alterations, functional impairments, and chronic pain. Given the dual vector’s effect on cartilage preservation, we tested if it could protect against OA-induced motor and pain-associated behavioral changes. CatWalk-based gait analysis showed significantly improved gait and motor functions in DMM mice receiving the dual vector compared to vehicle, as gait parameters such as paw print intensity, swing speed, and step pattern number were comparable to sham control mice (**Fig 1c,d, Suppl Fig S4d,e)**. Similarly, DMM-induced thermal hyperalgesia was attenuated by dual vector compared to vehicle-treatment, as measured by the hot plate assay (**Fig 1e**). Overall, these results show *HCAd-NFkb::IL-1Ra; EF1::Prg4* treatment attenuates injury-induced cartilage loss, alleviates pain-related behavioral changes, and improves locomotion in a PTOA mouse model.

### Dual vector treatment improves lameness and structural outcomes in a surgical large animal OA model

Pursuant to the murine dual vector gene therapy’s promising results in the DMM model, we investigated the treatment’s efficacy in a large animal model of OA (**Suppl Fig S1e,f**), leveraging the well-established equine carpal osteochondral fragment model (*12, 14, 36*). As before, we first confirmed equine dual vector-induced transgene expression in 293T cells in the presence or absence of TNFα-stimulation at the transcript (**Suppl Fig S3f,g**) and the protein level (**Suppl Fig S3h,i,k**). To induce OA, an osteochondral fragment was created in the distal radiocarpal bone in one randomly assigned middle carpal joint, while the contralateral joint underwent arthroscopic evaluation without fragmentation (sham). OA joints received either vehicle (PBS) or equine dual vector treatment while sham joints received an equal volume of PBS at 5 days post-surgery. Following recovery from surgery, animals began treadmill exercise (15min/session, 5 sessions/week for 2 months) to establish mild OA by 70 days (**Suppl Fig S1e,f, Suppl Table S1**). Horses receiving dual vector treatment showed improved lameness scores compared to vehicle controls, as subjectively graded using the American Association of Equine Practitioners (AAEP) lameness scale (**Fig 2a**), using a modified videographic lameness scale with the OA limb on the outside of a circle (**Fig 2b**), and objectively following flexion of the OA joint (**Fig 2c**). Of note, to account for inter-subject variability in surgical pain, lameness score changes relative to day 5 scores are shown. At termination of the observation period, horses receiving dual vector treatment showed improved articular cartilage structure and reduced T2 prolongation compared to vehicle controls, as measured by cartilage histologic scores and MRI-based analysis (**Fig 2d-e**). Prolonged T2 values are associated with disruption to the articular cartilage collagen network (*27, 28*), and MRI-based analysis revealed significantly less prolonged T2 mapping times for dual vector versus vehicle control groups for several locations both adjacent to (radiocarpal bone) and remote from the injury site (intermediate and 3rd carpal bones) (**Fig 2d, f**). Histological scoring of cartilage fibrillation of the 3^rd^ carpal bone was more severe in vehicle control treated joints compared to those treated with the dual vector (**Fig 2e, g, Suppl Fig S5**). Collectively, these data suggest *HCAd-NFkb::IL-1Ra; EF1::Prg4* effectively protects functional and structural whole joint integrity in a surgical equine OA model.

**Fig. 2.**
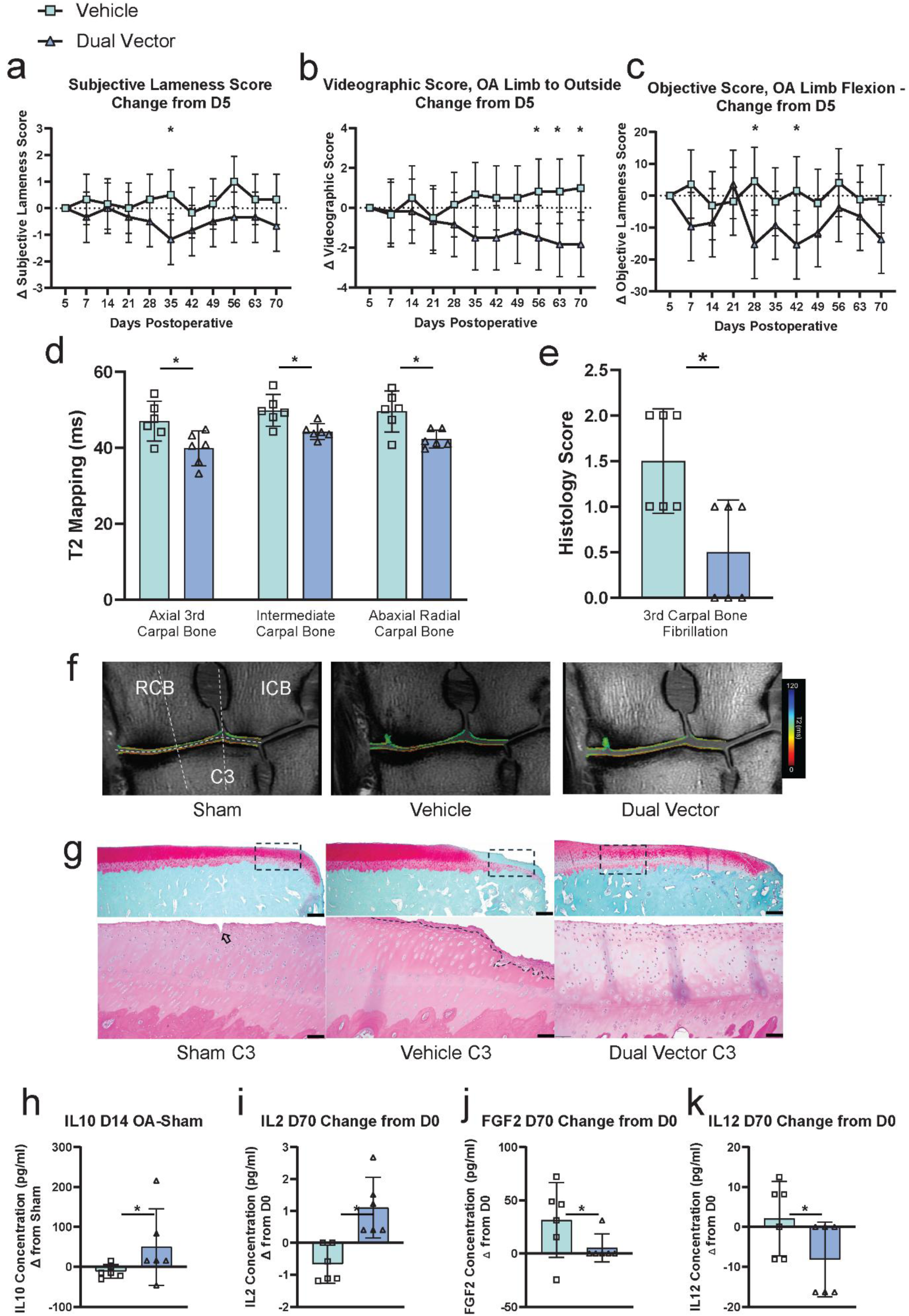
Efficacy of dual vector in equine OA. Results of the equine surgical OA model treated with *HCAd-NFkb::eqIL-1Ra; EF1::eqPrg4* (dual vector) or vehicle control. A) Change in subjective American Association of Equine Practitioners (AAEP) lameness score from day 5. B) Change in modified subjective videographic lameness score from day 5. C) Change in objective lameness score following flexion of the OA-affected limb from day 5. D) T2 mapping at the axial aspect of the 3^rd^ carpal bone, intermediate carpal bone, and abaxial aspect of the radial carpal bone measured by MRI. E) Histology scores of 3^rd^ carpal bone articular cartilage fibrillation. F) Representative MRI images from sham (left), vehicle treated (center), and dual vector treated (right) joints, showing T2 values (ms) of the articular cartilage surfaces. G) Representative safranin O/fast green (top images, scale bar 500μm) and hematoxylin/eosin (bottom images, scale bar 100μm) of 3^rd^ carpal bones from sham (left), vehicle treated (center), and dual vector treated (right) joints. Box inserts reflect area shown in the bottom images. Arrowhead indicates a focal site of fibrillation identified in a sham joint, while the dashed line shows an area of more extensive articular cartilage loss in a vehicle treated joint. * indicates p<0.05. Graphs display mean values, while error bars depict 95% confidence intervals for vehicle control (green squares); dual vector-treated joints (blue triangle).

### Cross-species attenuation of OA-linked joint inflammation following dual vector treatment

Inflammation is a key pathophysiological process in OA, and multiple cell and tissue types within the whole joint contribute to the chronic, low-grade inflammatory environment seen at various stages of disease pathogenesis (*17–20*). For example, macrophage infiltration into the arthritic joint capsule (synovium) is an accepted early indicator of OA, thought to contribute to structural changes and chronic pain development (*20*). To test the dual vector’s anti-inflammatory capacity across species, we first assessed whole joint macrophage infiltration in mouse knees using immunohistochemistry (F4/80; pan macrophage marker that labels M1- and M2-like populations). Dual vector treated OA joints contained significantly fewer F4/80-labeled macrophages compared to vehicle-treated OA joints (**Fig 3a,b; Suppl Fig S6**). Notably, while vehicle-treated OA joints contained significantly fewer anti-inflammatory CD206-labeled anti-inflammatory M2 macrophages, this reduction was normalized to levels detected in non-surgical controls (naïve, sham) in OA joints treated with *HCAd-NFkb::IL-1Ra; EF1::Prg4* (**Fig 3a,c; Suppl Fig S7**). These observations suggest that dual vector shifts macrophage polarization toward an anti-inflammatory M2-like state, supporting a protective immunomodulatory effect of this gene therapeutic approach in a PTOA mouse model.

**Fig. 3.**
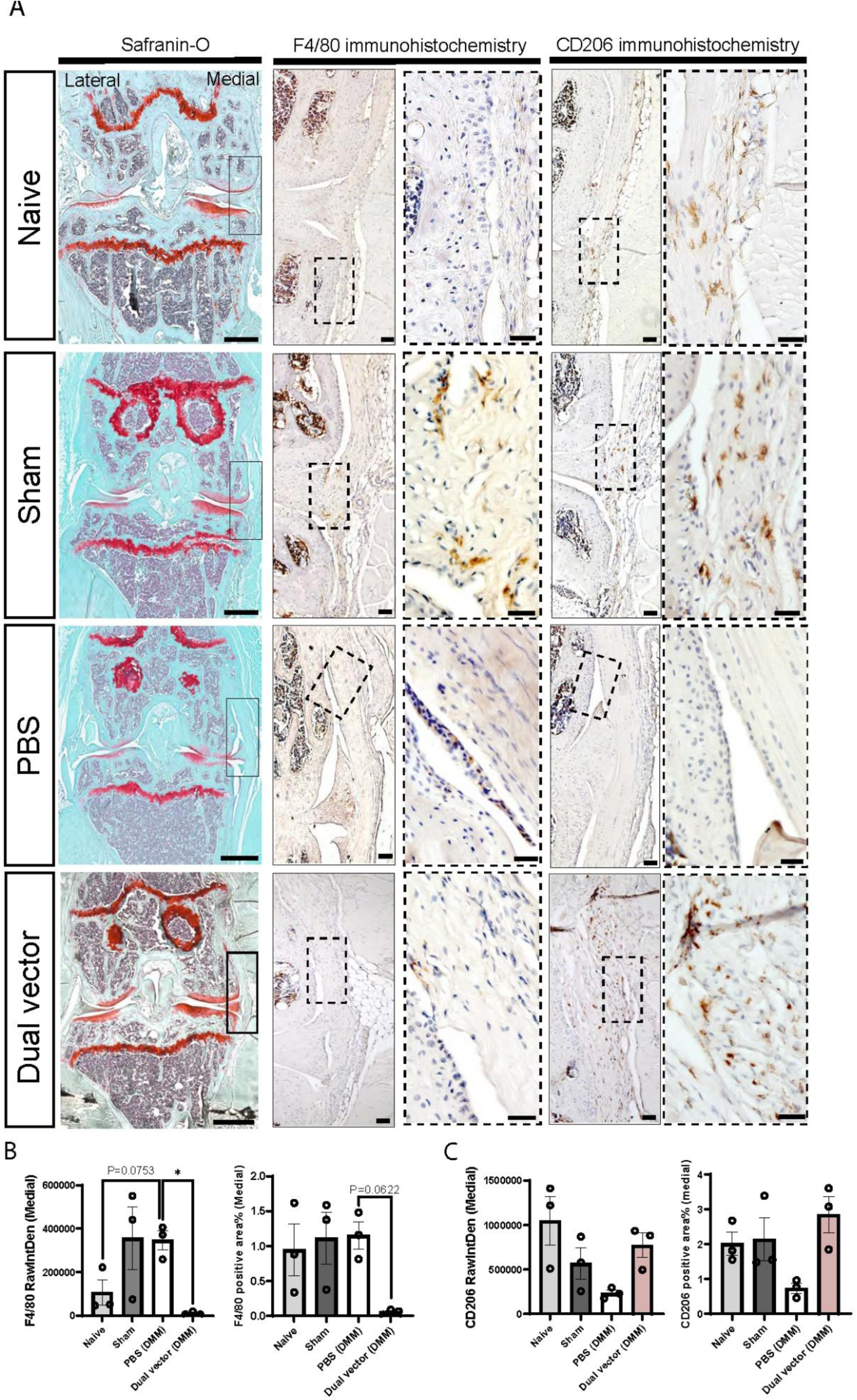
Dual vector treatment restores the macrophage landscape in the murine knee joint. Immunohistochemical assessment of F4/80-positive macrophage infiltration and CD206-associated M2-like macrophage polarization in mouse knee joints. (A) Representative immunohistochemistry images of F4/80 (pan-macrophage) and CD206 (M2-like macrophage) staining in mouse knee joint sections, shown at mid and high magnification alongside the corresponding low-magnification histology images (left column). Solid box-inserts in the low-magnification images indicate the regions shown at mid magnification, while dotted box-inserts in the mid-magnification images indicate areas shown at high magnification. (B) Quantification of integrated DAB-positive staining intensity (left) or DAB-positive area (right) within defined regions of interest (ROIs). Scale bars indicate 500 μm for Safranin-O staining images, 100 μm for low-magnification F4/80 and CD206 staining images, and 50 μm for high-magnification F4/80 and CD206 staining images. Statistical significance was assessed using Brown–Forsythe and Welch ANOVA with Dunnett’s T3 multiple-comparisons test. * = p<0.05. Mean values ±SEM are shown.

To determine how dual vector gene therapy modifies the inflammatory milieu in equine joints, we analyzed synovial fluid cytokine concentrations. Despite significant inter-animal variation, due to the outbred nature of the animals included in the study, dual vector-treated joints showed altered concentrations in several soluble factors within the synovial fluid compared to vehicle control joints (**Suppl Table S2,** note: many cytokines were below detection level). Synovial fluid IL-1Ra concentrations were significantly greater in dual vector treated joints on day 14 (**Suppl Fig S8a**), confirming inflammation-induced expression of this therapeutic target. Consistent with previous reports documenting a transient increase in lubricin/PRG4 expression in the first few days post injury (*37*), lubricin concentrations were greater in vehicle control treated joints compared to dual vector at day 5, followed by a trend for increased lubricin levels in dual vector-treated joints for the duration of the study, reaching significance at a late timepoint (day 42) (**Suppl Fig S8b, Suppl Table 3**). Dual vector-treated joints showed increased anti-inflammatory IL-10 concentrations in the OA limbs compared to vehicle-treated limbs when normalized to the contralateral sham joint at day 14 (**Suppl Fig S8c**). In addition, dual vector treatment increased synovial fluid IL-2, and moderately elevated IFNγ and fractalkine concentrations at endpoint measurements (day 70 vs day 0) compared to vehicle control (**Suppl Fig S8d-f**). Dual vector treated joints also showed reduced fibroblast growth factor 2 (FGF2) and IL-12 concentrations at endpoint measurements (day 70 vs day 0) compared to vehicle control (**Suppl Fig S8g,h**). While the effect of changes in immune cells and cytokines on tissue inflammation is typically challenging to interpret, the data suggest that joints treated with dual vector therapy showed increased concentrations of anti-inflammatory proteins and immunomodulatory cytokines, along with decreased concentrations of several pro-inflammatory cytokines and pro-catabolic factors compared to vehicle control joints.

### Dual vector treatment reduces OA-induced neuro-inflammation in the equine DRG

To elucidate potential chronic joint pain-mediating mechanisms at the level of the dorsal root ganglion (DRG) and determine the effect of dual therapy on these changes, we next performed single-nuclei RNA-seq (snRNA-seq) on bilateral equine eighth cervical (C8) DRGs from animals that received vehicle vs. dual vector treatment. Deep sequencing of 12 DRGs (sham vs OA, vehicle vs dual vector, n=3/group; 1.1 billion reads/sample) yielded 319,676 nuclei, with an average of 26,640 nuclei per sample (**Suppl Table S3**). Integrative analysis identified 20 major cell classes with specific marker gene expression profiles (**Fig 4a**,**Suppl Fig S9a-c**) that are largely consistent with those previously described in the cross-species DRG atlas (*38*), while offering higher resolution of immune cell populations than previously reported in other models due to the increased sequencing depth and cell number available in this large animal model. Despite the high degree of inter-subject variation expected in an outbred equine study population, comparing cell proportions in the placebo group (Sham vs OA side) revealed significant differences in DRGs that innervate affected (OA-side) compared to sham joints, including expansion of T cells, dendritic cells, and lymphatic endothelial cells, along with a reduction in neurons and repairing Schwann cells (**Fig 4b**). These findings complement previous chronic pain studies in a murine and human context (*39–45*). In contrast, dual vector-treated OA DRGs contained significantly fewer macrophages, mast cells, and capillary endothelial cells, while arachnoid fibroblasts were expanded compared to sham DRGs (**Fig 4c**). Direct comparison of cell type proportions in DRGs innervating dual vector-vs vehicle-treated OA joints revealed treatment-induced reductions in several glial and immune-related cell populations and expansion of vascular and fibroblast cell populations (**Suppl Fig S9d**). To overcome potential batch and inter-individual variation effects on our analysis, we next normalized OA-side cell proportions to those on the sham side, within each horse (i.e., [cell type 1 on OA side] / [cell type 1 on sham side]), and subsequently compared these normalized OA-linked cell populations between the vehicle and dual vector groups (**Fig 4d**; ratios ∼1 reflect similar proportions between sham and OA DRGs). Similar to the direct aggregated comparison (**Suppl Fig S9d**), dual vector treatment significantly reduced several immune cell populations (macrophages, T cells, mast cells), glial cells (satellite glia, non-myelinated Schwann cells), and lymphatic endothelial cells compared to the placebo group (**Fig 4d**). Interestingly, the IL-1 receptor (*IL1R1*) is most highly expressed in vascular cells and fibroblasts, whereas the endogenous IL-1 receptor antagonist (*IL1RN*) is predominantly expressed in macrophages (**Suppl Fig S9e**), suggesting the dual vector’s therapeutic effects may be relayed from the joint to the DRG through both vascular and neuronal pathways.

**Fig. 4.:**
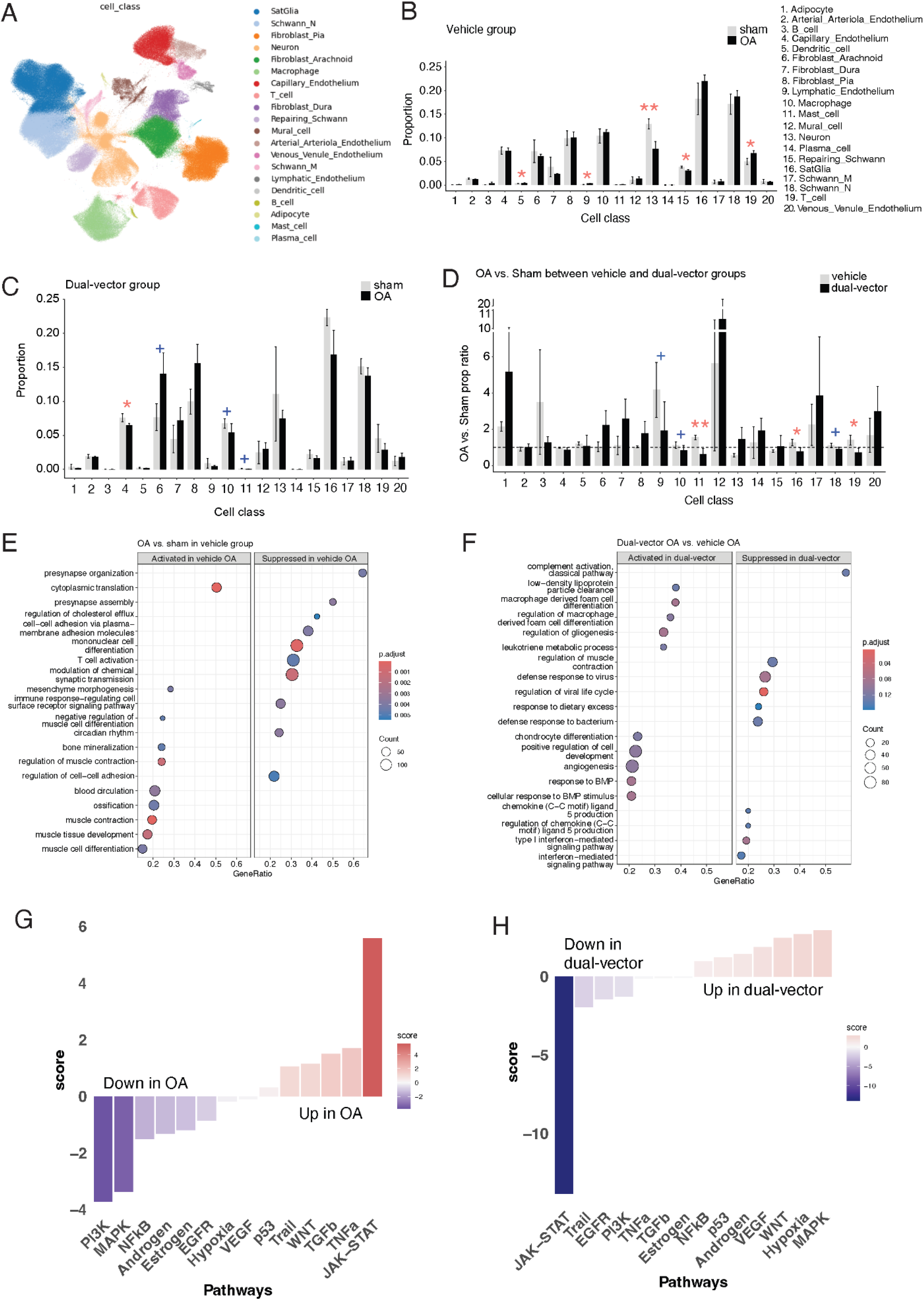
The effect of OA and gene therapy on equine DRGs at single cell resolution. (A) UMAP showing the transcriptomic cell atlas derived from snRNA-seq data from 12 horse DRGs, with cells colored by major cell class. (B) Bar plot showing the comparison of cell proportions between OA and Sham in the vehicle group. (C) Bar plot showing the comparison of cell proportions between OA and Sham in the dual-vector group. (D) Bar plot showing the normalized cell proportions in OA relative to sham between the placebo and dual-vector groups. Dotted line marks unaltered proportions for a given proportion in OA DRGs compared to sham ([cell type 1 in DRG on OA side] / [cell type 1 in DRG on sham side] = 1) (E) Dot plot showing the activated and suppressed biological processes in OA DRG compared with the sham DRG in the placebo group based on bulk RNAseq analysis. (F) Dot plot showing the activated and suppressed biological processes in OA DRG of the dual-vector group compared to the placebo group based on bulk RNAseq analysis. (G) Bar plot showing up- and down-regulated pathways in OA DRGs compared to sham controls in the placebo group. (H) Bar plot showing the up- and down-regulated pathways in dual-vector-treated OA DRGs compared to vehicle-treated DRGs. A one-tailed wilcoxon ranking sum test was conducted to calculate the P-values in Fig 4B-D. +: P≤0.1, *: P≤0.05, **: P≤0.01, and ***: P≤0.001. Bar graphs depict mean values ± standard errors.

Subsequent bulk RNA-seq of DRGs followed by differential gene expression (DEG; **Suppl Fig S9f,g**) and gene set enrichment analysis (GSEA) revealed that OA activates biological processes related to cytoplasmic translation, bone and muscle homeostasis, blood circulation, and others, whereas processes related to synaptic development and function, cholesterol homeostasis, cell-cell adhesion, and immune cell receptor signaling were suppressed in OA DRGs compared to sham in the vehicle group (**Fig 4e**). Comparing OA-induced changes between treatment groups revealed the dual vector activates several tissue repair processes, including macrophage differentiation, gliogenesis, chondrogenesis, response to BMP, and others. In contrast, gene therapy suppressed several inflammatory and immune response pathways in the OA DRG compared to OA-side DRGs from the vehicle group, including complement activation, viral defense responses, chemokine production, and type I interferon-mediated signaling (**Fig 4f**). Moreover, pathway enrichment analysis revealed strong OA-induced activation of JAK-STAT signaling (**Fig 4g**), a key immunomodulatory pathway involved in the acute-to-chronic pain transition through neuroinflammation and glial activation (*46–48*), while PI3K and MAPK pathways were down-regulated in the vehicle-treated group (**Fig 4g**). Importantly, *HCAd-NFκB::IL-1Ra-EF1::Prg4* strongly downregulated OA-induced JAK-STAT pathway activation compared to the placebo group (**Fig 4h**), further supporting the intra-articular dual vector’s effect on reducing neuroinflammation in the DRG.

### Dual vector treatment suppresses macrophage and T cell expansion in equine OA DRGs

To better characterize how joint disease and gene therapy alter neuro-immune interactions in the DRG, we further investigated cellular and molecular changes in two major immune cell classes: macrophages and T cells. Seven distinct macrophage populations were identified in the equine DRG, including five tissue-resident macrophage populations and two monocyte-derived macrophage populations (**Fig 5a**, 27,932 macrophages total, **Suppl Fig S10a-c**). Vehicle-treated OA DRGs revealed significantly expanded tissue-resident macrophages and moderate reduction of matrix-associated macrophages compared to contralateral sham DRGs (**Fig 5b**). In contrast, dual vector-treated OA DRGs contained markedly reduced macrophage populations compared to vehicle-treated DRGs, including tissue-resident macrophages and M1-like inflammatory macrophages when adjusting for inter-individual variation (**Fig 5c**; [cell type 1 on OA side]/[cell type 1 on sham side]). Notably, gene therapy-treated OA DRGs exhibited the lowest inflammatory macrophage signature across groups, as assessed by the M1/M2 macrophage ratio (**Fig 5d**). Subsequent pseudo-bulk analysis of vehicle-treated OA macrophages showed activation of cell proliferation-related processes and immune activation, along with suppression of biological processes involved in extracellular matrix organization, cell junction assembly, regulation of neuron projection development, glial cell differentiation, and cell surface receptor protein tyrosine kinase signaling pathway (**Fig 5e**). In contrast, dual-vector gene therapy reversed this signature, activating the processes that were suppressed in vehicle-treated OA DRGs along with additional repair processes (angiogenesis, ossification, and cell migration; **Fig 5f**). Pathway enrichment analysis revealed excessive inflammatory JAK-STAT signaling in vehicle-treated OA DRG macrophages, which was suppressed in dual vector-treated OA macrophages (**Fig 5g,h**). In contrast, proliferation and repair pathways (EGFR, TGFβ) were downregulated in vehicle-treated OA macrophages but significantly activated in dual vector-treated OA macrophages (**Fig 5g,h**).

**Fig. 5.:**
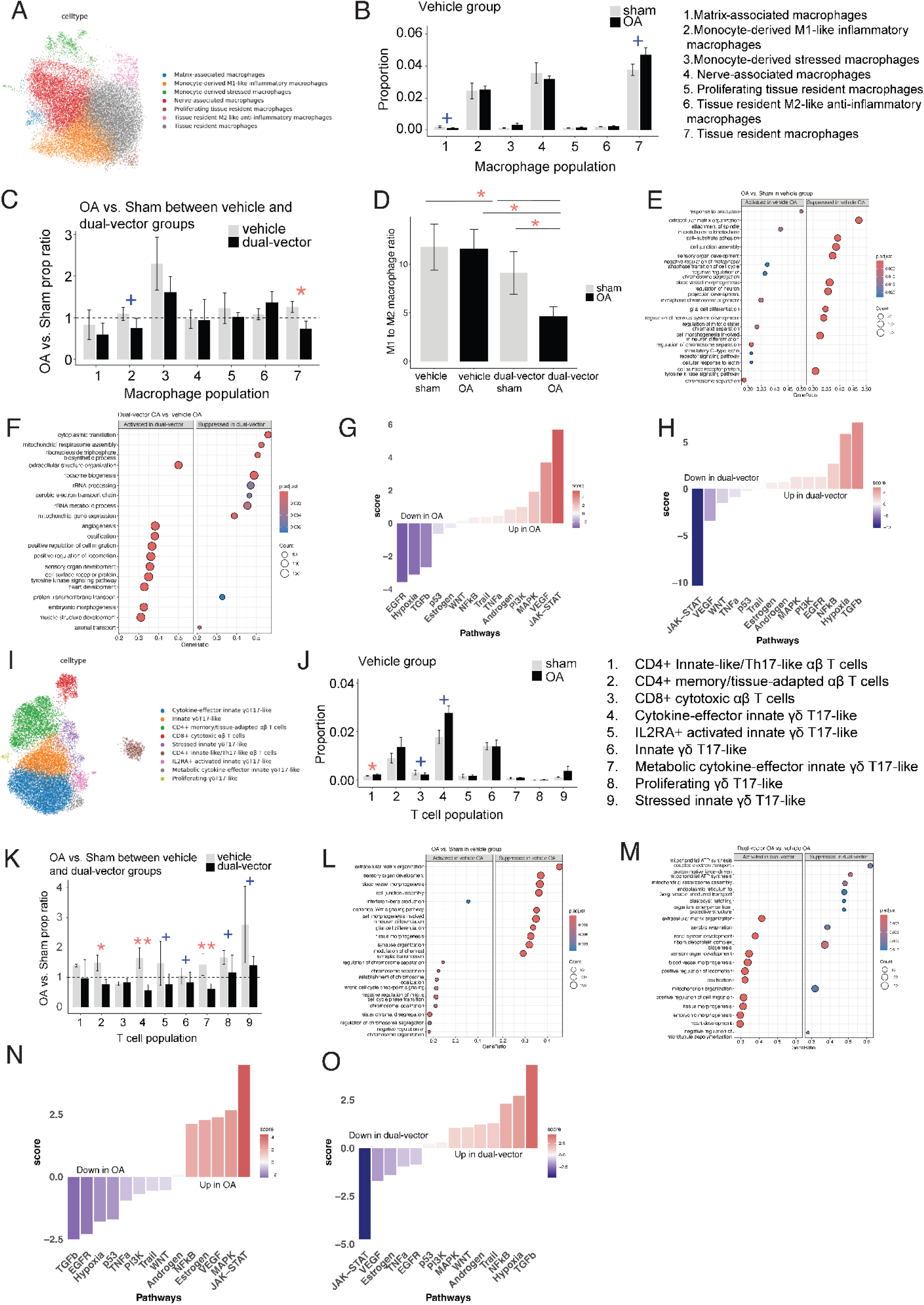
The effect of OA and gene therapy on equine macrophages and T cells in the DRG. (A) UMAP showing macrophages derived from snRNA-seq data from 12 horse DRGs, with cells colored by macrophage populations. (B) Bar plot showing the comparison of macrophage cell proportions between OA and Sham in the placebo group. (C) Bar plot showing normalized macrophage cell proportions in OA relative to sham between the placebo and dual-vector groups. Dotted line marks unaltered proportions for a given proportion in OA DRGs compared to sham ([cell type 1 in DRG on OA side] / [cell type 1 in DRG on sham side] = 1) (D) Bar plot showing the ratio of M1 vs. M2 macrophages for various groups. (E) Dot plot showing the activated and suppressed biological processes in OA macrophages compared with sham controls in the placebo group based on pseudo-bulk analysis. (F) Dot plot showing the activated and suppressed biological processes in OA macrophages from the dual-vector group relative to the placebo group based on pseudo-bulk analysis. (G) Bar plot showing the up- and down-regulated pathways in OA macrophages relative to sham controls in the placebo group. (H) Bar plot showing the up- and down-regulated pathways in OA macrophages treated with dual-vector compared to vehicle. (I) UMAP showing T-cells derived from snRNA-seq data from 12 horse DRGs, with cells colored by T cell populations. (J) Bar plot showing the comparison of T-cell proportions between OA and Sham in the placebo group. (K) Bar plot showing normalized T cell proportions in OA relative to sham between the placebo and dual-vector groups (L) Dot plot showing the activated and suppressed biological processes in OA T cells relative to sham controls in the placebo group based on pseudo-bulk analysis. (M) Dot plot showing the activated and suppressed biological processes in OA T cells from the dual-vector group relative to the placebo group based on pseudo-bulk analysis. (N) Bar plot showing the up- and down-regulated pathways in OA T-cells relative to sham controls in the vehicle-treated group. (O) Bar plot showing the up-regulated and down-regulated pathways in OA T cells treated by dual-vector relative to the vehicle group. A one-tailed wilcoxon ranking sum test was conducted to calculate the p-values in Fig 5B, C, D, J, and K. +: P≤0.1, *: P≤0.05, **: P≤0.01, and ***: P≤0.001. Bar graphs depict mean values ± standard errors.

T-cell subtype analysis revealed nine transcriptionally distinct populations, including six *γδ* Th17-like and three infiltrating αβ T-cell populations (**Fig 5i**, 14,437 T-cells total). Compared to sham controls, vehicle-treated OA DRGs exhibited an expansion of CD4^+^ Innate-like/Th17-like αβ T-cells, along with trends to increased cytokine-effector innate *γδ* 17-like T-cells and reduced CD8^+^ αβ T-cells (**Fig 5j**). Mirroring the macrophage response, dual-vector gene therapy normalized OA-induced changes when adjusting for inter-individual variation (**Fig 5k**). Consistent with the observed T-cell expansion phenotype, pseudo-bulk analysis of vehicle-treated OA T-cells revealed activation of cell proliferation-related processes and IFNβ production, along with suppression of biological processes involved in extracellular matrix organization, cell junction assembly, neuronal and glial differentiation, and canonical Wnt signaling (**Fig 5L**). The dual vector gene therapy activated some of the biological processes that were suppressed in vehicle-treated DRGs, similar to what was observed in macrophages (**Fig 5M**). Consistently, JAK-STAT signaling was up-regulated and TGFβ signaling was reduced in vehicle-treated OA DRG T-cells (**Fig 5N**), whereas gene therapy down-regulated reversed this signature (**Fig 5O**). Collectively, these results suggest that intra-articular dual vector treatment effectively attenuates immune activation and boosts tissue repair in OA DRGs.

### Exploring the neurobiological mechanisms of chronic joint pain in the equine DRG

We next examined OA- and treatment-induced molecular and cellular changes in the sensory neuronal subtypes in the DRGs to explore the molecular and cellular mechanisms that may mediate chronic joint pain. The equine DRG contains 16 transcriptionally distinct neuronal populations, including two A-fiber proprioceptor populations, four A-fiber populations that may interact with other cell classes, three A*β*-low-threshold mechanoreceptors (LTMRs), one A*δ*-LTMR, two C-fiber non-canonical peptidergic (NP) nociceptors, three C-fiber peptidergic (PEP) nociceptors, and one C-fiber temperature-sensitive (C-Thermo) population (**Fig 6A; Suppl Fig S11a-c**). Interestingly, OA was associated with significant reductions of multiple subtypes, covering several A-, A*β*-LTMR- and C-fiber populations (**Fig 6b**), complementing previous clinical studies of chronic pain conditions (*43, 45*). Only two subtypes, A*δ*-LTMRs and thermosensitive C fibers, trended towards expansion in vehicle-treated OA DRGs. Remarkably, dual vector treatment reversed nearly all OA-linked neural population changes when adjusting for inter-individual variation, expanding populations that showed OA-linked neural loss while reducing populations that had expanded in the presence of joint disease (**Fig 6c**).

**Fig 6.**
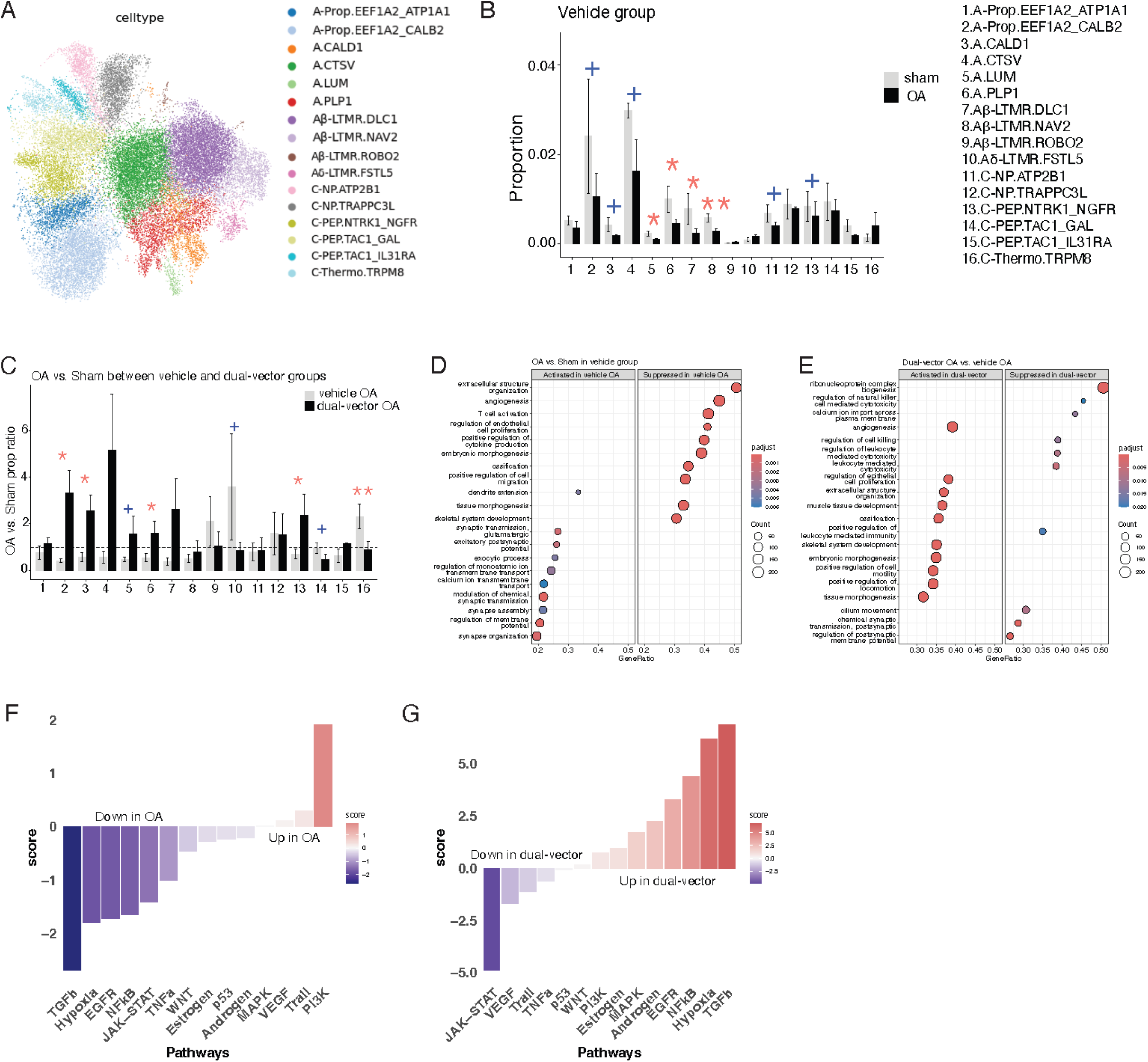
Neurons in DRGs of equine models of OA and therapeutic intervention. (A) UMAP showing neurons derived from snRNA-seq data from 12 horse DRGs, with cells colored by neuronal populations. (B) Bar plot showing the direct comparison of neuronal cell proportions between OA and sham in the vehicle group. (C) Bar plot showing normalized neuronal cell proportions in OA relative to sham between the placebo and dual-vector groups. Dotted line marks unaltered proportions for a given proportion in OA DRGs compared to sham ([cell type 1 in DRG on OA side] / [cell type 1 in DRG on sham side] = 1). (D) Dot plot showing the activated and suppressed biological processes in OA neurons compared with sham controls in the placebo group based on pseudo-bulk analysis. (E) Dot plot showing the activated and suppressed biological processes in OA neurons from the dual-vector group relative to the placebo group, based on pseudo-bulk analysis. (F) Bar plot showing the up- and down-regulated pathways in OA neurons relative to sham controls in the vehicle-treated group. (G) Bar plot showing the up- and down-regulated pathways in OA neurons treated by dual-vector relative to the vehicle group. A one-tailed wilcoxon ranking sum test was conducted to calculate the P-values in Fig 5B and C. +: P≤0.1, *: P≤0.05, **: P≤0.01, and ***: P≤0.001. Bar graphs depict mean values ± standard errors.

To probe OA-associated changes in biological processes and pathways that govern sensory neural biology and function, pseudo-bulk analysis of vehicle-treated OA sensory neurons revealed activation of numerous biological processes related to neuronal excitability compared to sham sensory neurons (**Fig 6d; Suppl Fig S11d,e**). This data suggests that OA sensory neurons may reside in a state of hyperexcitability, an important part of the peripheral sensitization process that occurs during the transition from acute to chronic pain (*49*). Notably, dual vector treatment suppressed several neural activity-related processes, as well as pathways involved in leukocyte mediated cytotoxicity and immune response, while activating several tissue repair processes (**Fig 6e**). Subsequent pathway enrichment analysis revealed activation of PI3K signaling, previously associated with pain and inflammation, and downregulation of tissue-repair-associated TGFβ signaling in vehicle-treated OA DRGs (**Fig 6f**). In contrast, sensory neurons in dual vector-treated OA DRGs activated TGFβ signaling while suppressing the inflammatory JAK-STAT pathway (**Fig 6g**).

## DISCUSSION

Osteoarthritis (OA) is a complex, progressive disease that affects the whole joint as an organ and, due to a lack of disease-modifying therapies, remains a primary cause of chronic pain and disability worldwide (*1, 50*). Despite coordinated efforts, no FDA-approved treatment exists that effectively slows OA progression and sustainably reduces chronic joint pain. Standard-of-care treatment for OA-related pain includes non-steroidal anti-inflammatory drugs (NSAIDs), steroids, or opioids, which fail to provide long-term joint pain relief and can pose significant health and dependency risks (*51–53*). The etiopathology of OA is complex, with distinct yet interconnected processes and pathways contributing to degeneration of joint tissues. In a recent paradigm-shift, research shows that OA represents a spectrum of conditions with partially overlapping clinical (phenotype) and molecular (endotype) manifestations and underlying pathophysiology (*5, 54–56*). Hence, developing effective treatments for OA will likely require strategies that simultaneously target multifactorial drivers of joint degeneration, inflammation, and pain. Gene therapy can overcome many of the practical issues that have challenged OA therapeutic development, given its ability to balance both sustained and regulated target gene expression via localized delivery-mechanisms that minimize systemic or off-target effects (*57*). High-capacity adenoviral (HCAd)-based gene therapeutic platforms have several features that make them the vector of choice for developing affordable, multi-factorial therapeutic approaches for common, complex diseases such as OA, cancer, and others (*57–59*). First, HCAd’s large payload capacity (∼36kb) enables generation of single vectors carrying multiple therapeutic candidates (*27*). This property is particularly relevant for use and combination of large therapeutic genes, such as *Prg4*. Second, intra-articular HCAd delivery confers high transduction rates of joint-associated tissues (*27, 28*). Third, HCAds are economical to produce at scale due to high yields and scalable production methods. Building on this premise, several studies have shown encouraging results, including amelioration of pain and OA progression using HCAd-vectors in small and large animal models (*14, 26–28, 35*). Phase 2 clinical studies with *HCAd-NFκB::IL-1Ra* are underway (NCT06884865), with phase 1 data suggesting long-term pain (>2 years) reductions in OA patients after a single intra-articular injection (*29–31*). In addition, intra-articular delivery of 2 HCAd vectors that separately express IL-1Ra and Prg4 provided improved protection in a murine OA model compared to either single vector alone (*35*). However, the feasibility and efficacy of combinatorial HCAd-based treatment for OA, using a bicistronic vector harboring two therapeutic candidates, has not been explored to date.

Here, we developed a combinatorial gene therapy consisting of an HCAd-based vector carrying two therapeutic transgenes (IL-1Ra and PRG4) and investigated its efficacy in a mouse, rat, and equine OA model. Exploratory efficacy analysis in the rat ACLT model showed enhanced protection of the dual vector compared to vectors encoding individual therapeutic targets, confirming our previous study(*35*). In addition, intra-articular delivery of this dual cistron vector (*HCAd-NFκB::IL-1Ra;EF1::Prg4*; dual vector) improved structural, functional, and pain outcomes in a murine and equine OA model (**Fig 1,2**), and reduced cellular (mouse, horse) and molecular (horse) markers of inflammation in the affected joint (**Fig 3**). Importantly, multi-modal transcriptomic profiling of equine DRGs that contain joint-innervating sensory neurons reveals extensive neuro-inflammation and molecular signatures of neural hyperexcitability in the context of joint disease, which was largely restored by gene therapy (**Fig 4-6**). Collectively, these findings suggest that intra-articular dual vector treatment reduces OA progression in the joint, reduces the inflammatory environment in the joint and the DRG, and reduces known mechanisms of peripheral sensitization in the DRG while decreasing pain metrics in small and large preclinical OA models.

There are several limitations of this study. First, we assessed the effect of gene therapy or vehicle on OA compared to sham-operated joints, in which the joint capsule is opened and closed without further injury. While this represents the conventional approach for surgical disease models, sham surgery causes transient post-surgical pain in rodents (*60, 61*). Hence, sham joints and DRGs do not represent a neutral baseline and may contain elevated inflammation and pain mechanisms compared to naïve animals. Second, this study contains limited measures of pain that are difficult to uncouple from functional joint impairment (hot plate and catwalk for mice; lameness for horses). Studying pain in preclinical models is challenging, as animals have developed strategies to hide pain-related behaviors and as pain behaviors are highly susceptible to external factors (*62*). While sophisticated machine-learning methods are being developed to overcome these challenges in murine and equine OA models, they were not available at the initiation of this study. Thus, further studies leveraging these cutting-edge approaches will be needed to better correlate how molecular findings in DRGs of chronic pain models correlate with pain-related behaviors. Third, the greater genetic variability between individual horses compared to more traditional, inbred preclinical models, such as mice, can add additional heterogeneity to the response to surgical induction of OA, severity of OA, gene expression, and treatment responses. Despite this variability, treatment with the dual vector improved several functional, structural, and molecular outcome measures; the clear improvements noted in the equine OA model suggest high translatability to human OA.

The equine carpal osteochondral fragment model has been used with some variation for several decades (*13*). Traditionally, this OA model utilizes the American Association of Equine Practitioners (AAEP) lameness grading scale. While this scale enables standardized clinical lameness grading, it has a limited dynamic range, especially for mild lameness. Complementing AAEP lameness grades with a modified videographic scoring system allowed for further discernment of lameness at a trot in a straight line and on circles using a 0-10 scale (**Fig 2b**). In addition, we included objective measures alongside all subjective scoring, including the use of an inertial sensor system in grading lameness (**Fig 2c**). Objective lameness scoring is increasingly being used to evaluate lameness in the carpal osteochondral fragment model (*63–65*). Limitations of the inertial sensor-based system in this and past studies include the inability to detect bilateral forelimb lameness and difficulties interpreting natural asymmetries which are present when horses travel on a circle (*66*). Combining subjective and objective measurements for lameness parameters broadened the potential to capture more subtle treatment differences given the relatively mild degree of lameness. Inter-horse variability in surgical inflammation and discomfort is an additional source of inconsistency, which was recently corrected for by assigning treatment groups only following ranking of postoperative lameness scores (*64*). While this current study’s design was not amenable to a similar strategy, we instead normalized to postoperative, pre-treatment lameness scores obtained on day 5 to account for inter-subject variability in surgical pain. Finally, while magnetic resonance imaging (MRI) in this model has been described (*65, 67*), this is the first time T2 mapping times are documented for the equine carpal osteochondral fragment model, revealing a significant difference in prolongation, with less prolonged values seen with dual vector therapy compared to vehicle control, indicative of sustained normal articular cartilage collagen structure in treated joints.

Transcriptomic studies of equine DRGs are limited to a single spatial transcriptomic proof-of-concept study (*68*), whereas bulk or single nucleus/cell RNAseq-studies have so far focused on other equine tissues. This study presents the first multi-modal transcriptomic reference dataset for the equine DRG, combining deep RNAseq (1.1 billion reads/sample) with deep single nucleus RNAseq (a median UMI count of 4641.7/cell). The resulting single cell reference atlas of the equine DRG contains 20 major cell classes (**Fig 4**) that can be further separated into multiple subtypes (**Fig 5-6**). Equine major cell classes are largely consistent with those recently described by the PRECISION network’s cross-species DRG atlas, including human, mouse rat, guinea pig, rhesus macaque and cynomolgus macaque DRGs (*38*). Additionally, we expanded the annotation of fibroblast, endothelial, and immune cell populations in the DRG.

Cellular and molecular analyses of major DRG cell types, and of macrophages, T-cells, and sensory neurons equine DRGs reveal an inflammatory cellular environment with highly activated JAK-STAT signaling (**Fig 4-6**), confirming previous reports in mouse and human models of chronic pain (*39–44*). Both comparisons revealed OA-induced expansion of cytokine-effector innate γ*δ* T-17-like cells, and trending increases in several other γ*δ* T-subtypes (**Fig 5j**). γ*δ* T-cells produce the pro-inflammatory cytokine IL-17 and often localize in close proximity to sensory neurons in the periphery, where they influence neuro-immune crosstalk (*69, 70*). A recent study reported γδ^+^ T-cell-expansion in murine OA joints, increasing IL-17 levels that promoted disease progression (*71*). However, few studies have explored γ*δ* T-cell function in pain pathways: while acute inflammatory pain-related metrics seem unaltered in mice lacking γ*δ* T-cells (*72*), these cells’ role in other pain types remains unstudied (*73*). Thus, further studies are needed to elucidate how γ*δ* T-cells alter pain experiences, and whether they play a role in chronic pain. Importantly, intra-articular gene therapy reversed many of the observed OA-linked population changes. Of note, comparisons between cell type proportions changes induced by OA or gene therapy were conducted using two parallel analyses: direct comparison of a cell type’s proportion between both groups (**Suppl Fig 8d**) versus a comparison following inter-individual combination for the proportion observed in the contralateral sham DRG (**Fig 4d,5c,5k, 6c**). Both analyses have limitations: while direct comparison is unable to compensate for inter-individual variability, the normalized comparison may overcompensate due to gene therapy’s potential effect on the contralateral sham side. However, their strong correlation support that the “true values” are expected to fall somewhere within both analyses.

Notably, OA induced significant loss of several sensory neural subtypes in equine DRGs (**Fig 6b**), a phenomenon recently reported in human DRGs from patients with neuropathic pain (*43, 45*). OA pain is complex, containing nociceptive, inflammatory, and neuropathic components that occur to varying degrees in a patient- and time-specific manner (*74*). Our observations suggest a neuropathic component to the pain experience of equine osteochondral fragment patients, confirming the validity and relevance of this model for better understanding the etiopathological mechanisms of OA. Previous studies have shown that synovial joints are primarily innervated by small C-fiber nociceptors, with additional contributions by nociceptive A*δ*-fibers, low-threshold mechanoreceptors (LTMRs), and proprioceptors (*75–81*). While these studies were performed in murine, leporine, and human joints, it is reasonable to assume that similar neural subtypes innervate the equine carpal joint. In addition, joint innervation was found to be dynamic, with excessive branching of CGRP- or Nav1.8-expressing nociceptive neuronal fibers (C-fibers, A*δ*-fibers, LTMR) in OA joints (*80–82*). The observed proportional changes in many of these neural subtypes and the reversal –or perhaps even overcompensation–of this phenotype by intra-articular gene therapy (**Fig 6d**) suggest chronic joint disease may induce joint-innervating sensory neuronal loss, a hypothesis that should be further explored. Importantly, equine sensory neurons showed strong activation of processes related to neuronal excitability, a well-documented phenomenon during the acute-to-chronic pain transition (*44, 83*), and these processes were suppressed by *HCAd-NFκB::IL-1RA;EF1::Prg4*. Together, our transcriptomic analyses of the equine DRG identify potential molecular and cellular bio-signatures of correction for chronic joint pain that may be translatable to other pain types and interventions.

In sum, these results show that dual vector-treatment confers generalized protection from OA progression, reduction of joint inflammation, and pain reduction in two preclinical OA models with distinct anatomical and functional joint characteristics. In addition, dual vector-treatment partially suppresses OA-mediated neuroinflammation, neuronal loss, and molecular mechanisms of neural hyperexcitability at the level of the DRG, revealing potential correction mechanisms and bio-cellular signatures of chronic joint pain. Building on previous experiences with the *HCAd-NFκB::IL-1Ra* mono-therapy, which showed similar cross-species efficacy and is currently in phase 2 clinical trials, the dual vector is a prime candidate for testing in the clinical setting, with the potential for greater effect than monotherapy given its dual design targeting both joint health and inflammation.

## Materials and Methods

This description represents an abbreviated version of the materials and methods. More details can be found in the supplementary materials. All researchers involved in data collection, data processing, and statistical analysis remained blinded to group allocation throughout the study.

### Vector Design and in vitro Validation

All dual vector constructs encoding mouse, rat, or equine *NFkB::IL-1RA* and *EF1::PRG4* were packaged into a high-capacity adenoviral vector (HCAd). For in vitro validation of murine and equine dual vector constructs, 293 cells (human embryonic kidney cells) were transduced with *HCAd0* (empty vector control), or with HCAd’s encoding a *LacZ* control, murine or equine *NFkB::IL-1RA* or *EF1-PRG4* single, and *NFkB::IL-1RA;EF1::PRG4* dual vectors. To assess IL-1Ra transcript and protein levels, 293 cells were incubated with recombinant human TNF-α (25 ng/ml), 24 hr after transduction with NF-κB-IL-1RA or dual vector to induce NF-κB regulated IL-1RA expression. 48 hr after transduction, the conditioned medium was collected for ELISA and quantitative real-time PCR analysis.

### OA Models and Treatment

#### Rats

9- to 10-week-old Sprague Dawley rats (n=15), acquired from Envigo and housed at the Inotiv, Inc, Boulder, CO under their IACUC protocol, were subjected to ACLT surgery, in which the anterior cruciate ligament (ACL) was transected through an incision of the joint capsule, upon which animals were randomly assigned to 4 treatment groups. Seven days later, 20 µL of vector formulation buffer (vehicle control), or 5×10^8^ viral particles of *HCAd-NFkb::rIL-1Ra* or of *HCAd-rEF1::Prg4* (single gene vectors) or of *HCAd-NFkb::rIL-1Ra; EF1::rPrg4* (dual gene vector; dual vector) was injected intra-articularly. Fifteen weeks later, rats were euthanized, and knee joints were harvested.

#### Mice

FVB/N mice were obtained from the Center for Comparative Medicine (CCM), Baylor College of Medicine (BCM) in Houston, Texas. All animal procedures were approved by the BCM Institutional Animal Care and Use Committee (IACUC). Twelve-week-old male FVB/N mice were randomly assigned to undergo bilateral destabilization of the medial meniscus (DMM) or sham surgery as previously described (*8, 28, 35*). Briefly, the medial meniscotibial ligament was transected to destabilize the medial meniscus, inducing OA. Sham-operated mice underwent identical joint exposure without ligament transection. Animals that received DMM surgery were randomly allocated into control or treatment groups, each receiving bilateral intra-articular HCAd injections. Controls received 5 μl PBS (vehicle control), while experimental animals received *HCAd-NFkb-mIL-1RA::EF1-mPRG4* (murine dual vector; 1×10^8^ viral particles per joint) 1 week post surgery. Sham-operated mice were not injected.

#### Horses

Twelve skeletally mature light breed horses were enrolled in the study with institutional animal care and use approval (**Suppl Table S1;** IACUC 2022-0014), and the study was performed under the Association for Assessment and Accreditation of Laboratory Animal Care (AALAC)’s guidelines. Prior to surgery, horses were conditioned to treadmill exercise, which was resumed 14 days following surgery 5 times weekly. Treadmill sessions consisted of walking (5-6km/h) for 3 minutes, trotting (13-14km/h) for 3 minutes, cantering (25-28km/h) for 3 minutes, trotting for 3 minutes, and walking for 3 minutes, for a total exercise time of 15 minutes. OA was induced using a previously described distal radiocarpal bone osteochondral fragment model under general anesthesia (*14, 36*). Briefly, an 8-mm osteochondral fragment was created in the distal radiocarpal bone of one randomly selected middle carpal joint, followed by burring of the underlying subchondral bone to create a ∼5-mm defect. Horses were stall-rested and bandaged until suture removal. Horses were then randomly allocated to *HCAd-NFkb-eqIL-1RA::EF1-eqPRG4* (equine dual vector; 2×10^11^ viral particles) or vehicle (saline). On day 5 post-operatively, synovial fluid was aspirated from each joint, and OA joints were injected with the appropriate treatment diluted to a total volume of 3 mL in PBS. Sham-operated joints received an injection of 3 mL PBS.

### Murine Motor and Pain Behavior Assessments

Thermal sensitivity was determined using the hot plate assay, as previously described (*35*). Gait analysis was performed using the CatWalk XT system (Noldus Information Technology) in BCM’s Neurobehavior Core according to the core’s protocol, described in supplementary methods. Several parameters commonly used for translational osteoarthritis studies, (maximum contact maximum intensity mean, swing time mean, number of step-sequence patterns, total step number) were analyzed and reported (*84*).

### Equine Outcome Measures

All clinical assessments were performed by the same board-certified large animal surgeon, who was blinded to treatment group allocation. Weekly clinical assessments included subjective and objective lameness evaluation, including response to flexion and subjective evaluation of joint effusion and range of motion. Lameness examinations were recorded and assessed on a straight line and circles in both directions. Subjective lameness examination followed American Association of Equine Practitioners (AAEP) 0-5 grading. Objective lameness evaluation was performed using an inertial sensor system (Equinosis Q Lameness Locator^®^) (*63*). Flexion tests were performed on both carpi, with subjective grading of the flexed limb on a 0 (no response) to 3 (severe lameness following flexion) scale in addition to objective scoring. Joint effusion was scored subjectively on a 0 (no palpable or visible effusion) to 3 (severe effusion which cannot be indented with palpation of the dorsal joint pouch) scale and objectively using carpal circumference measurements at the bottom of the accessory carpal bone, which was analyzed using the ratio to baseline measurement to account for variation in carpal size. (**Table S3**).

Baseline clinical assessment was performed within 5 days of surgery, followed by day 5 after surgery and prior to injection, day 7 following surgery, then weekly through day 70. Baseline synovial fluid was obtained from both middle carpal joints at the time of arthroscopy and prior to saline distention, at day 5 immediately prior to injection, at day 14 following surgery and weekly through day 70.

On days 0, 14, and 70, synovial fluid cell counts and differentials were performed on EDTA-treated synovial fluid. Following euthanasia with pentobarbital sodium at day 70, both carpi underwent MRI, were grossly evaluated and scored by a single observer using a published scoring system (*85*). Synovial and osteochondral tissues were collected for histology. Bilateral dorsal root ganglia (DRG’s) from the C6-T2 spinal nerves were collected, flash frozen in liquid nitrogen, and stored at -80°C.

### Equine Synovial Fluid analysis

Synovial Lubricin and IL-1Ra concentrations were measured using enzyme linked immunosorbent assays (ELISA) (*36*). For a more comprehensive cytokine analysis, synovial fluid samples were thawed and treated with hyaluronidase (Streptomyces hyalurolyticus, 1U/ml in PBS, Millipore Sigma) for 30min at 37°C prior to performing the MILLIPLEX^®^ Equine Cytokine/Chemokine Panel (Millipore Sigma). All Luminex assays were performed at UC Davis’ Biologic and Molecular Analysis Core at UC Davis.

### Histological analysis

Upon collection, rat knee joints were decalcified for 7 days, trimmed, bisected in the frontal plane, and both halves were formalin-fixed, paraffin embedded in a single block cut side down. Blocks were sectioned (8 µm), mounted on glass slides, stained with toluidine blue using standard methods, fixed and processed, and paraffin-embedded, prior to being histologically processed. Sections were stained with toluidine blue and scored by a blinded pathologist according to the OARSI scoring system, as previously described (*86*). To minimize the high experimental variability induced by the surgery in this model, histological analysis was performed on the subset of animals that similarly responded to the ACLT surgery, as determined by their fibrosis score as a surrogate for instability development and weight bearing on the injured limb post-surgery.

Mouse knee joints were harvested 12 weeks post-surgery and fixed in 4% paraformaldehyde for 48 hr at 4 °C, washed three times in PBS, and decalcified in 10% EDTA at 4 °C for up to 10 days. Tissues were dehydrated, paraffin-embedded and sectioned at 6 μm. Joint sections were stained with Safranin O/Fast Green to assess structural damage of cartilage and proteoglycan content. Histologic sections were graded by Osteoarthritis Research Society International (OARSI) scoring (*86*). To study macrophages in mouse knees, antibodies against F4/80 (70076S, Cell Signaling Technology; dilution - 1:200) and CD206 (24595S, Cell Signaling Technology; dilution - 1:200) were used, as previously described (*87*).

Equine synovial membrane tissues were fixed in 2% formalin for 5-7 days and processed for paraffin embedding. Osteochondral sections were obtained from the radial, intermediate, and third carpal bones using an oscillating saw. Osteochondral sections were fixed in 2% formalin for 5-7 days, decalcified using 40% formic acid/20% sodium citrate decalcification solution, and processed and paraffin embedded. Tissue sections (5μm) were stained with hematoxylin/eosin (synovial membrane and osteochondral) or Safranin O/Fast Green (osteochondral) and graded by a board-certified veterinary pathologist using previously described grading criteria (*13*).

### Phase-contrast microCT Analysis

For 3D cartilage volumetric analysis of mouse knees, left hind limbs were processed as previously described (*88*), and scanned by Bruker SkyScan 1272 scanner in BCM’s Optical Imaging and Vital Microscopy Core.

### Single Nuclei RNA Sequencing (snRNAseq) of equine DRGs

Snap-frozen equine DRGs were processed using the GentleMACS Octo Dissociator with C-tubes (Miltenyi Biotec) in ice-cold Nuclei Extraction Buffer according to the manufacturer’s protocol (see suppl methods). Nuclei concentrations were determined using a hemocytometer, and nuclei were visually inspected under fluorescence microscopy to assess nuclear integrity. Single-nucleus libraries were generated using the Chromium Single Cell 3′ Gene Expression v4 platform (10x Genomics), targeting approximately 20,000 nuclei per library. Libraries were sequenced on both Illumina and Ultima platforms at the BCM sequencing core to increase sequencing depth (90). Sequencing reads from both platforms were jointly processed using Cell Ranger Count (v9.0.1)(*89*), and mapped to the EquCab3 reference genome with Ensemble v114 annotation. Quality control was performed using a public pipeline (91), including ambient RNA correction with SoupX (*90*), doublet removal with DoubletFinder (v2.0.4) (92), and low-quality cell filtering (UMI count < 500, gene count < 500, or > 5% of mitochondrial percentages).

Data integration and clustering were performed using the scVI pipeline (scVI-tools v1.2.0 and Scanpy v1.10.3, with sample ID as the batch variable, n_layer=2, n_latent=30, and 2000 to 10,000 highly variable genes). Leiden algorithm-based clustering (resolutions 0.1-1)was used to determine the optimal number of clusters. Major cell class annotation and sub-clustering of macrophages, T cells, and neurons was performed using the same scVI pipeline, based on established marker genes. Cell type proportions were calculated as the fraction of total cells per DRG sample. Cell proportion differences across conditions were assessed using a one-tailed Wilcoxon rank-sum test.

### Bulk RNA-seq Data Processing

Raw Ultima sequencing reads from 12 equine DRGs were preprocessed with a custom script to generate single-end reads compatible with bulk RNA-seq analyses, then aligned to the EquCab3 genome with Ensemble v114 annotation using STAR (2.7.11b) (*91, 92*).

### Differential Gene Expression Analyses, Gene Set Enrichment and Pathway Enrichment

For snRNA-seq data, UMI counts were aggregated at the sample level by cell class to generate pseudo-bulk profiles, with three replicates per surgery type per treatment group. Differentially expressed genes (DEG) analysis was performed using edgeR (v4.4.0) and variancePartition (v1.36.2)(*93, 94*)., after filtering lowly expressed genes (mean CPM ≤0.5). OA effects were assessed by comparing OA DRGs with contralateral sham DRGs per cell class, using the model: ∼ surgery_type + (1 | animal_ID). Gene therapy effects were evaluated by comparing dual-vector treated OA DRGs with vehicle-treated OA DRGs, using the model: ∼treatment_group.

For bulk RNA-seq, gene-level read counts were generated using htseq-count package (v2.1.2) (*95*). DEG analysis for OA- and gene therapy-associated effects were performed using the same edgeR (v4.4.0) and variancePartition (v1.36.2) pipeline applied to the snRNA-seq pseudo-bulk data (*93, 94*).

Gene set enrichment analysis was performed using gseGO from clusterProfiler (v4.14.6) (*96*). with genome-wide log2 fold changes from DEG analyses as input and the GO Biological Process ontology as the reference library. Enrichment significance was assessed using 100,000 permutations, with multiple-testing correction performed using the Benjamini-Hochberg method. Gene set and pathway enrichment analyses were performed on DEG results from bulk RNA-seq and pseudo-bulk snRNA-seq data respectively.

### Statistical Analysis

All clinical and synovial fluid data were analyzed using mixed linear models including fixed effects of treatment, day, and treatment*day interaction, with horse included as a random effect. Data evaluated only at study termination, including gross joint, MRI, and histologic scoring, was evaluated for normality by visual observation of Q-Q plots and a Shapiro-Wilk test. Normally distributed data was evaluated using a Welch’s t-test, while non-parametric data was evaluated using a Mann-Whitney test. Murine data were analyzed using Brown–Forsythe and Welch ANOVA followed by Dunnett’s T3 multiple-comparisons test. For *in vitro* studies comparing multiple groups, ordinary one-way ANOVA with multiple-comparisons test were used. Statistical significance was considered α< 0.05. Statistical analyses were performed using GraphPad Prism and RStudio.

## List of Supplementary Materials

Materials and Methods

Supplementary figures: Fig. S1 to Fig S11

Supplementary tables: Table S1 to Table S3

Additional references, cited in supplementary methods

## Acknowledgments

This work was supported by the U.S. Department of Defense Congressionally Directed Medical Research Programs (CDMRP) (W81XWH-22-1-0372), the Lawrence Family Bone Disease Program of Texas, by NICHD’S IDDRC (P50HD103555 to BCM’s Rodent Neurobehavior Core), and by the RE-JOIN consortium (*97*), an NIH HEAL-funded team science initiative to identify the biological underpinnings of chronic joint pain (UC2AR082200).

We acknowledge BCM’s Optical Imaging & Vital Microscopy Core (OIVM) for assistance with phase-contrast µCT imaging, BCM’s rodent neurobehavior core for support with murine pain behavioral analysis, BCM’s single cell core for snRNAseq support, and UC Davis’ Biologic and Molecular Analysis Core for performing Luminex assays.

## Author contributions

Conceptualization: ES, JW, ZD, JY, KG, YB, NAH, HR, BL

Methodology: ES, JW, ZD, JY, CM, MW, JAA, DC, RC, JBE, DP, MJ, CL, IY, SP, LY, SV

Investigation: ES, JW, ZD, JY, CM, MW, JAA, DC, RC, SP, JBE, DP, MJ, CL, IY, SP, LY, SV

Visualization: ES, JW, ZD, JY, CM, JAA, SP, JBE, MJ, CL, LY, SV

Funding acquisition: PN, TD, KCW, KG, YB, NAH, HR, BL

Project administration: YB, NAH, HR, BL

Supervision: YB, NAH, HR, BL

Writing – original draft: ES, JW, ZD, YB, NAH, HR, BL

Writing – review & editing: ES, JW, ZD, JY, CM, MW, JAA, DC, RC, JBE, DP, MJ, CL, IY, SP, LY, SV, RSR, JDW, BRA, RC, KCW, RE-JOIN, PN, KG, YB, NAH, HR, BL

## Data, code, and materials availability

All data, code, and materials used in the analysis will be made available through RE-JOIN’s HEAL-approved data repository (SPARC) in some form to any researcher for purposes of reproducing or extending the analysis. Viral vectors generated through this study will be shared through standard materials transfer agreements (MTAs).

## Other acknowledgments

We are grateful to members of the RE-JOIN consortium, an NIH HEAL-funded team science initiative to identify the biological underpinnings of chronic joint pain, for their suggestions and comments on this work. The RE-JOIN consortium consists of: Armen Akopian, Kyle Allen, Alejandro Almarza, Benjamin Arenkiel, Yangjin Bae, Bruna Balbino de Paula, Anita Bandrowski, Mario Danilo Boada, Jacqueline Boccanfuso, Jyl Boline, Dawen Cai, Dellina Carpio, Robert Caudle, Racel Cela, Yong Chen, Rui Chen, Brian Constantinescu, Cortez, Ibdanelo, Yenisel Cruz-Almeida, M. Franklin Dolwick, Chris Donnelly, Zelong Dou, Joshua Emrick, Malin Ernberg, Danielle Freburg-Hoffmeister, Spencer Fullam, Janak Gaire, Akash Gandhi, Benjamin Goolsby, Stacey Greene, Nele Haelterman, Michael Iadarola, Shingo Ishihara, Azeez Ishola, Sudhish Jayachandran, Zixue Jin, Frank Ko, Priya Kulkarni, Zhao Lai, Brendan Lee, Yona Levites, Carolina Leynes, Jun Li, Martin Lotz, Lindsey Macpherson, Tristan Maerz, Camilla Majano, Anne-Marie Malfait, Maryann Martone, Bella Mehta, Richard Miller, Rachel Miller, Michael Newton, Alia Obeidat, Merissa Olmer, Dana Orange, Miguel Otero, Kevin Otto, Folly Patterson, Marlena Pela, Sienna Perry, Theodore Price, Hernan Prieto, Russell Ray, Dongjun Ren, Margarete Ribeiro Dasilva, Alexus Roberts, Elizabeth Ronan, Oscar Ruiz, Shad Smith, Mairobys Socorro, Kaitlin Southern, Joshua Stover, Michael Strinden, Hannah Swahn, Evelyne Tantry, Sue Tappan, Luis Tovias Sanchez, Airam Vivanco-Estela, Joost Wagenaar, Lai Wang, Kim Worley, Joshua Wythe, and Jiansen Yan.

## Supplementary figures

**Fig. S1.**
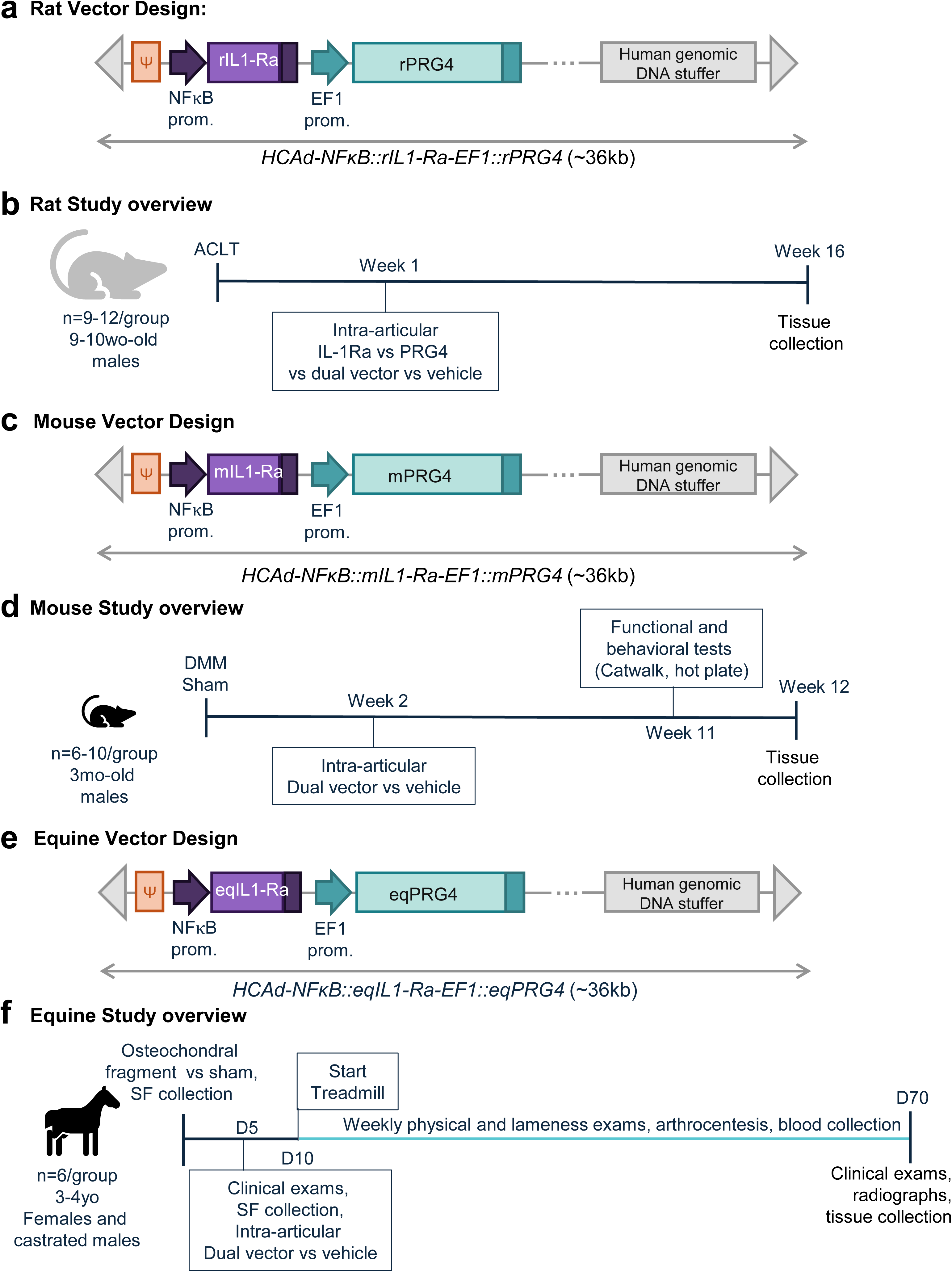
Study overview. (A) Overview of the rat dual vector (*HCAd-NFκB::rIL-1RA-EF1::rPrg4*) (B) Rat study design. 9- to 10-week-old male mice (n=9-12/group) were subjected to anterior cruciate ligament (ACLT) surgery and received intra-articular injections with HCAd vectors encoding individual therapeutic genes (IL-1Ra or PRG4) or dual vector 1-week post-surgery. Knee joints were collected for histopathological scoring at 16 weeks post-surgery. (C) Overview of the murine dual vector (*HCAd-NFκB::mIL-1RA-EF1::mPrg4*) (D) Mouse study design. 3-month-old male mice (n=6-10/group) were subjected to destabilization of the medial meniscus (DMM) or sham surgery and received intra-articular dual vector or vehicle injections. Functional and behavioral measures (hot plate, catwalk) were obtained 11 weeks post-surgery and tissues were collected at 12 weeks post-surgery. (E) Overview of the equine dual vector (*HCAd-NFκB::eqIL-1RA-EF1::eqPrg4*) (F) Equine study design. 3- to 4-year old castrated male and female animals (n=6/group) were subjected to osteochondral fragment or sham surgery. 5 days post-surgery, clinical exams were performed, synovial fluid (SF) was collected, and animals received intra-articular dual vector or vehicle treatment. Animals started a regular exercise regimen (15min/session, 5 sessions / week for 2 months) and received weekly physical and lameness exams, arthrocentesis, synovial fluid and blood collection until termination at day 70, when clinical exams, radiographs, and tissues were collected.

**Supplemental Fig. S2.**
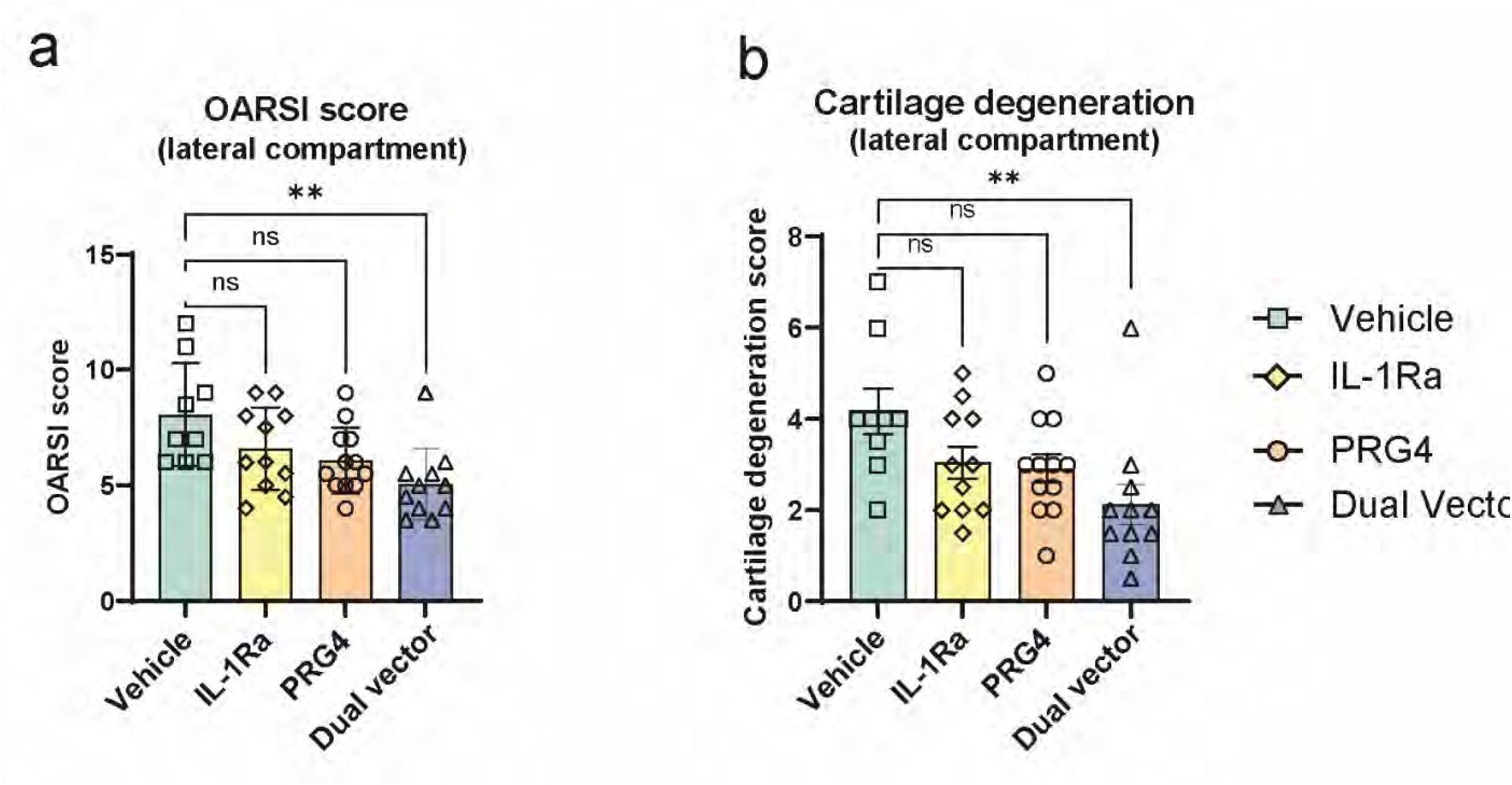
Results of the rat ACLT OA model treated with vehicle control (vector formulation buffer), with single-gene-encoding HCAds (*HCAd-NFkb::rIL-1Ra* or *HCAd-EF1::rPrg4*), or with *HCAd-NFkb::rIL-1Ra; EF1::rPrg4* (dual vector). (A) Composite OARSI scores for the lateral compartment of the joint, summed for each animal, based on Toluidine Blue-stained sections from vehicle-treated OA joints (vehicle, n=9), mono-treatment (IL-1Ra or PRG4; n=11 or n=12 respectively), or dual-vector treated OA joints (n=10). (B) OARSI scoring of articular cartilage degeneration in the lateral compartment of the joint for the same samples shown in A. Statistical significance was assessed using a Kruskal-Wallis test combined with Dunn’s multiple-comparisons testing. ** = p<0.01. Graphs depict mean values ±SD.)

**Fig. S3.**
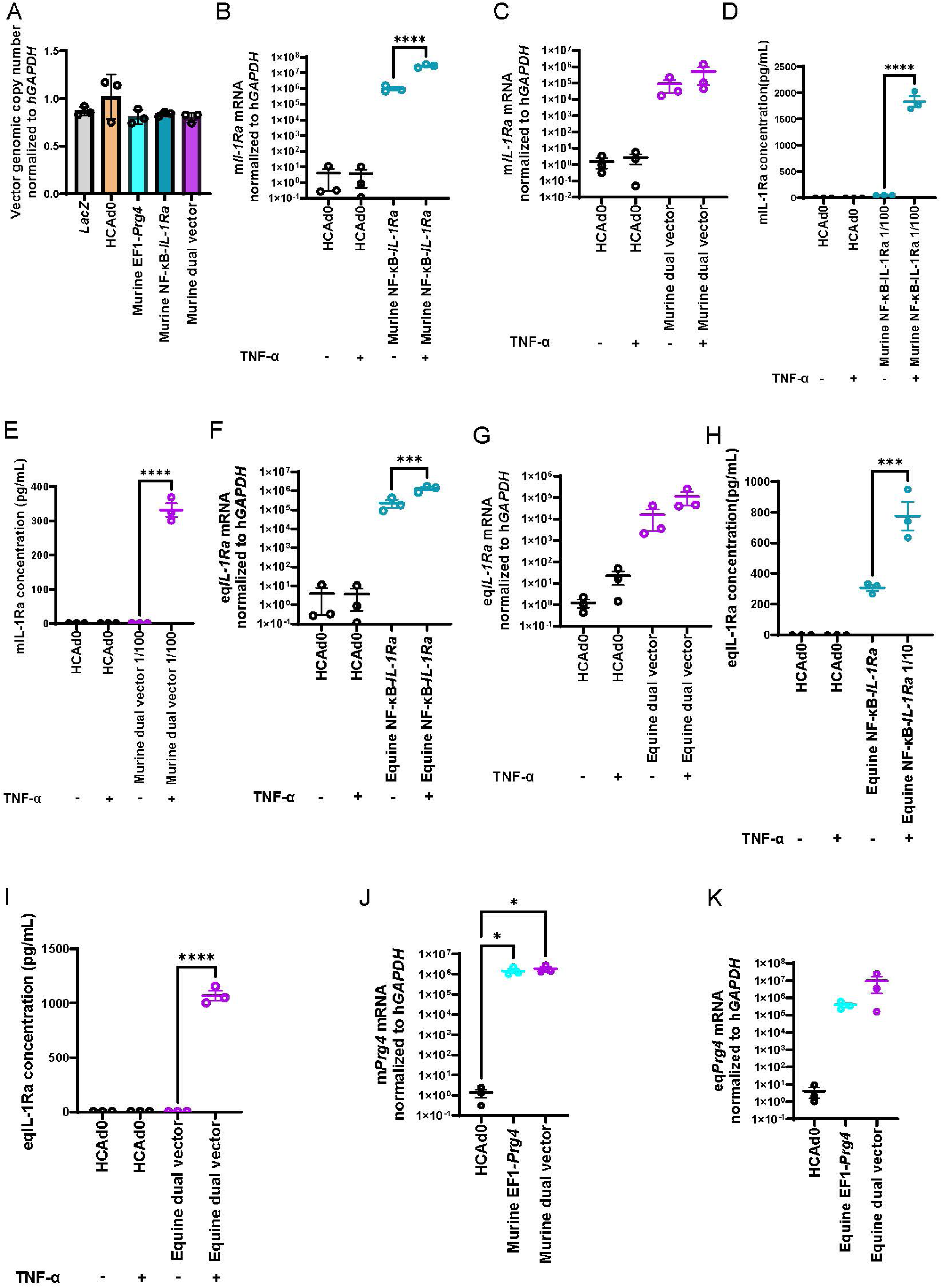
*In vitro* functional validation of HCAd vectors. (A) Genomic copy number of HCAd vectors expressing the indicated target genes. (B and C) Relative murine *Il1ra* transcript levels in HEK293 cells transduced with murine IL-1Ra or dual-expression vectors, with or without TNFα stimulation. (D and E) Murine IL-1Ra protein expression levels in HEK293 cells transduced with murine IL-1Ra or dual-expression vectors, with or without TNFα stimulation. (F and G) Relative equine *IL1RA* transcript levels in HEK293 cells transduced with equine IL-1Ra or dual-expression vectors, with or without TNFα stimulation. (H and I) Equine IL-1Ra protein expression levels in HEK293 cells transduced with equine IL-1Ra or dual-expression vectors, with or without TNFα stimulation. (J and K) Relative murine and equine *Prg4/PRG4* transcript levels in HEK293 cells transduced with murine or equine PRG4 and dual-expression vectors, respectively. Statistical significance was assessed using ordinary one-way ANOVA with Tukey’s multiple-comparisons test in (A), (J), and (K), and ordinary one-way ANOVA with Šídák’s multiple-comparisons test in (B)–(I). * = p<0.05, *** = p<0.001, **** = p<0.0001. Graphs show mean values ±SEM. Type or paste caption here. Create a page break and paste in the figure above the caption.

**Fig. S4.**
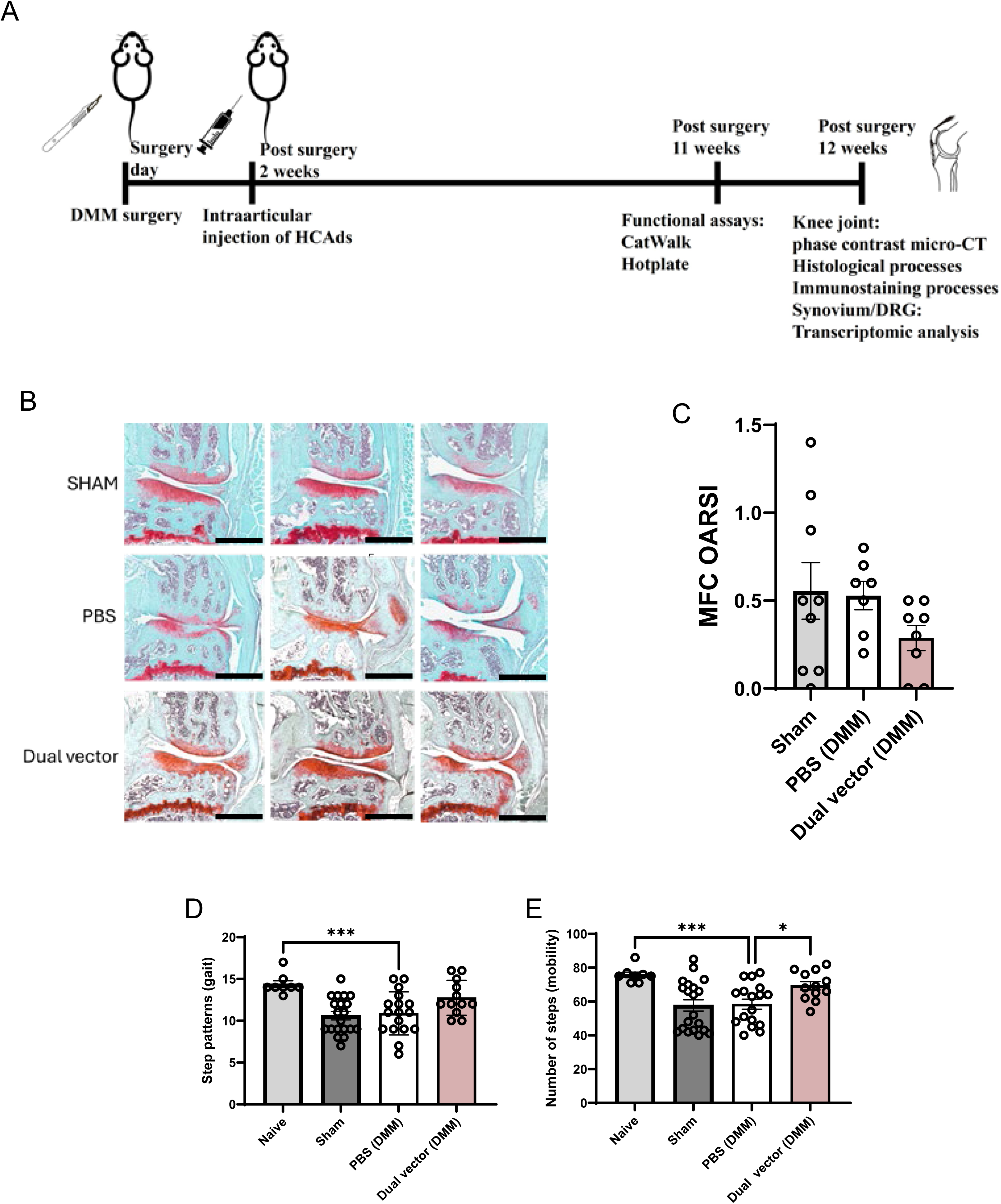
Mouse study overview, knee joint histology, and additional functional assessments. (A) Schematic overview of the mouse experimental timeline and procedures, including DMM surgery, intra-articular injection, downstream functional testing, and structural and histological analyses. (B) Representative coronal sections of Sham, vehicle (PBS)-treated, and dual-vector– treated mouse knee joints stained with Safranin O and Fast Green staining. (C) OARSI scoring of articular cartilage degeneration based on Safranin O-stained knee joint sections from Sham, vehicle-treated OA joints (PBS), and dual-vector treated OA joints. (D) CatWalk-based quantification of step patterns. (E) CatWalk-based quantification of mouse mobility by number of steps. Scale bars indicate 500 μm for Safranin-O staining images. Statistical significance was assessed using Brown–Forsythe and Welch ANOVA with Dunnett’s T3 multiple-comparisons test. * = p<0.05, *** = p<0.001. Graphs depict mean values ±SEM.

**Fig. S5.**
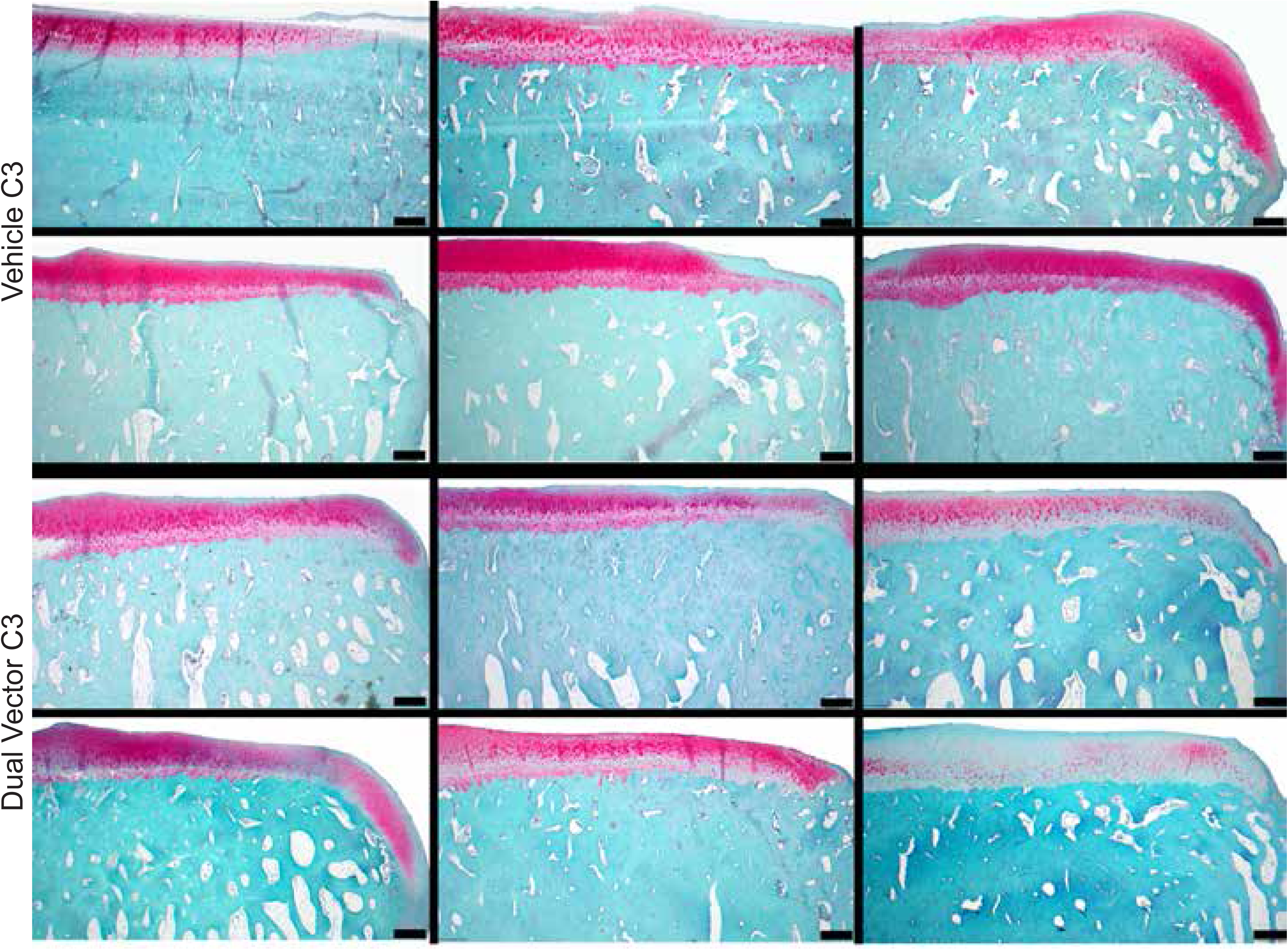
Equine joint histology. Composite images of safranin O/fast green (scale bar 500μm) of all 3^rd^ carpal bones from vehicle treated joints (top 2 rows) and dual vector treated joints (bottom 2 rows).

**Fig. S6.**
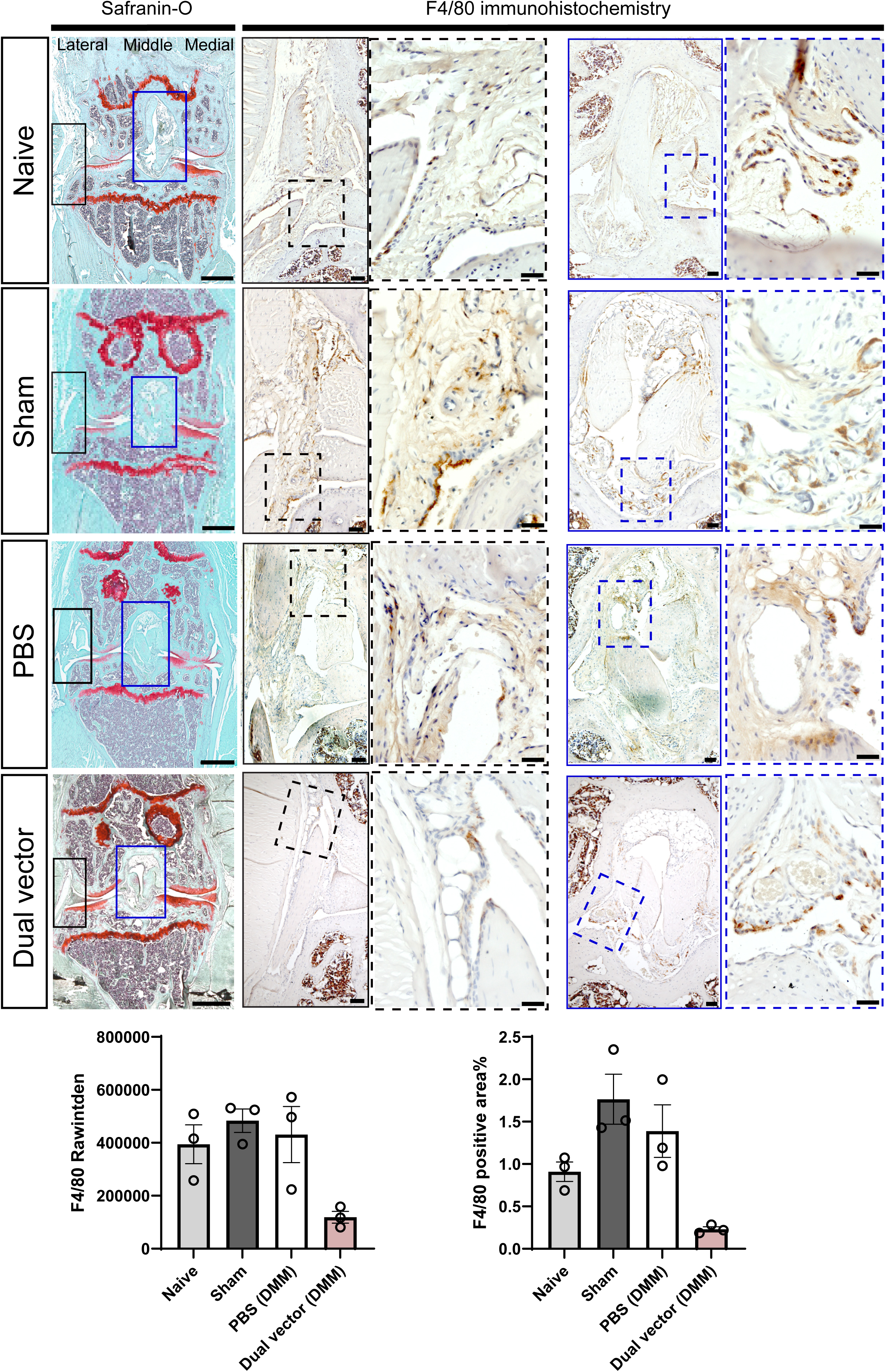
Immunohistochemistry images of F4/80-labeled macrophages, focusing on lateral and middle regions of mouse knee joints. (A) Representative F4/80 immunohistochemistry images, labeling all macrophage populations, are shown at medium and high magnification, with corresponding low-magnification histological images provided for anatomical orientation. Black and red boxes in the low-magnification histology images indicate the lateral and medial regions, respectively, analyzed at medium magnification. Black and red dotted boxes in the medium-magnification images indicate the corresponding lateral and medial regions shown at high magnification. (B) Integrated DAB-positive staining intensity and the corresponding percentage of DAB-positive area were quantified within defined ROIs. For each sample, measurements from the lateral, middle, and medial regions were averaged to generate a single value for analysis. Scale bars indicate 500 μm for low-magnification histology images, 100 μm for medium-magnification immunohistochemistry images, and 50 μm for high-magnification immunohistochemistry images. Statistical significance was assessed using Brown–Forsythe and Welch ANOVA with Dunnett’s T3 multiple-comparisons test. Graphs depict mean values ±SEM.

**Fig. S7.**
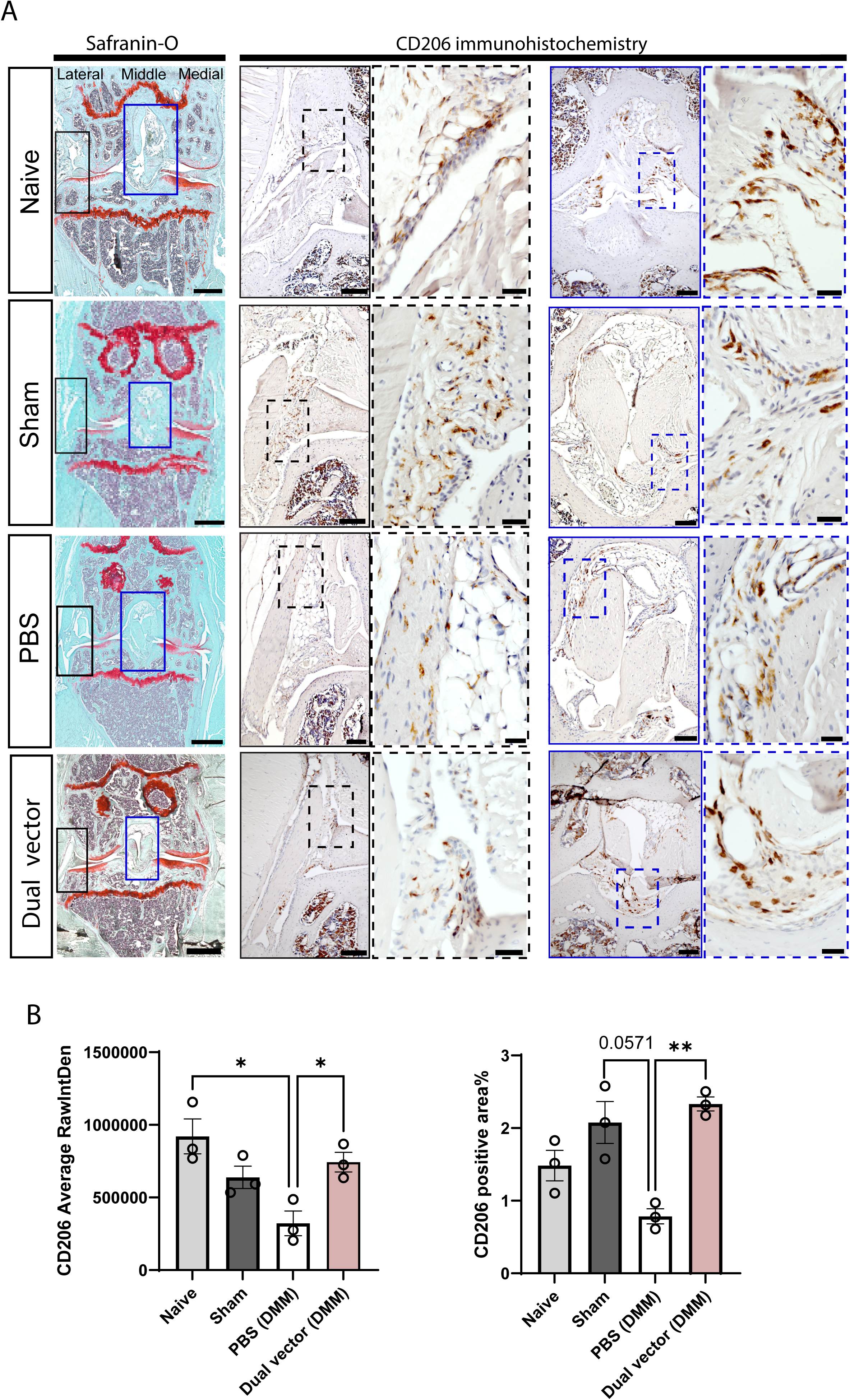
Immunohistochemistry images of CD206-labeled M2 macrophages, focusing on lateral and middle regions of mouse knee joints. (A) Representative CD206 immunohistochemistry images are shown at medium and high magnification, with corresponding low-magnification histological images provided for anatomical orientation. Black and red boxes in the low-magnification histology images indicate the lateral and medial regions, respectively, analyzed at medium magnification. Black and red dotted boxes in the medium-magnification images indicate the corresponding lateral and medial regions shown at high magnification. (B) Integrated DAB-positive staining intensity and the corresponding percentage of DAB-positive area were quantified within defined ROIs. For each sample, measurements from the lateral, middle, and medial regions were averaged to generate a single value for analysis. Scale bars indicate 500 μm for low-magnification histology images, 100 μm for medium-magnification immunohistochemistry images, and 50 μm for high-magnification immunohistochemistry images. Statistical significance was assessed using Brown–Forsythe and Welch ANOVA with Dunnett’s T3 multiple-comparisons test. * = p<0.05, ** = p<0.01. Graphs depict mean values ±SEM.

**Fig. S8.**
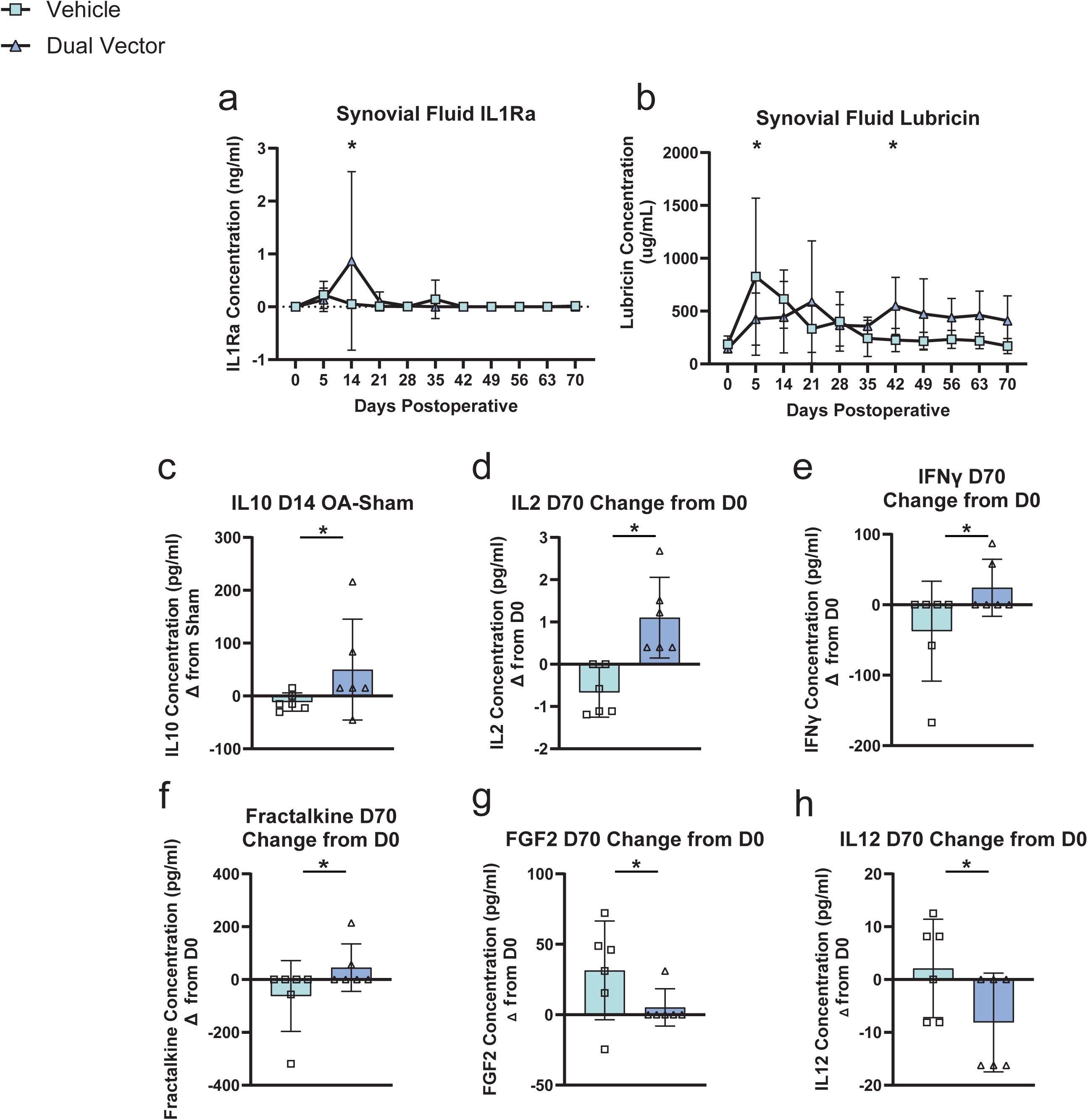
Synovial fluid soluble marker analysis in equine joints. Additional results of the equine surgical OA model treated with *HCAd-NFkb::IL-1Ra; EF1::Prg4* (dual vector) or vehicle control. A) Synovial fluid IL-1Ra concentrations. B) Synovial fluid lubricin concentrations. C) Synovial fluid IL-10 concentrations shown as the difference between OA limb and sham limb on day 14. D) Synovial fluid IL2 concentrations shown as the difference between day 70 and day 0 in the OA joints. E) Synovial fluid IFNγ concentrations shown as the difference between day 70 and day 0 in the OA joints. F) Synovial fluid fractalkine concentrations shown as the difference between day 70 and day 0 in the OA joints. G) Synovial fluid FGF2 concentrations shown as the difference between day 70 and day 0 in the OA joints. H) Synovial fluid IL12 concentrations shown as the difference between day 70 and day 0 in the OA joints. Graphs display mean values ± 95% confidence intervals for vehicle control (green squares); dual vector-treated joints (blue triangle). * indicates p<0.05.

**Fig. S9.**
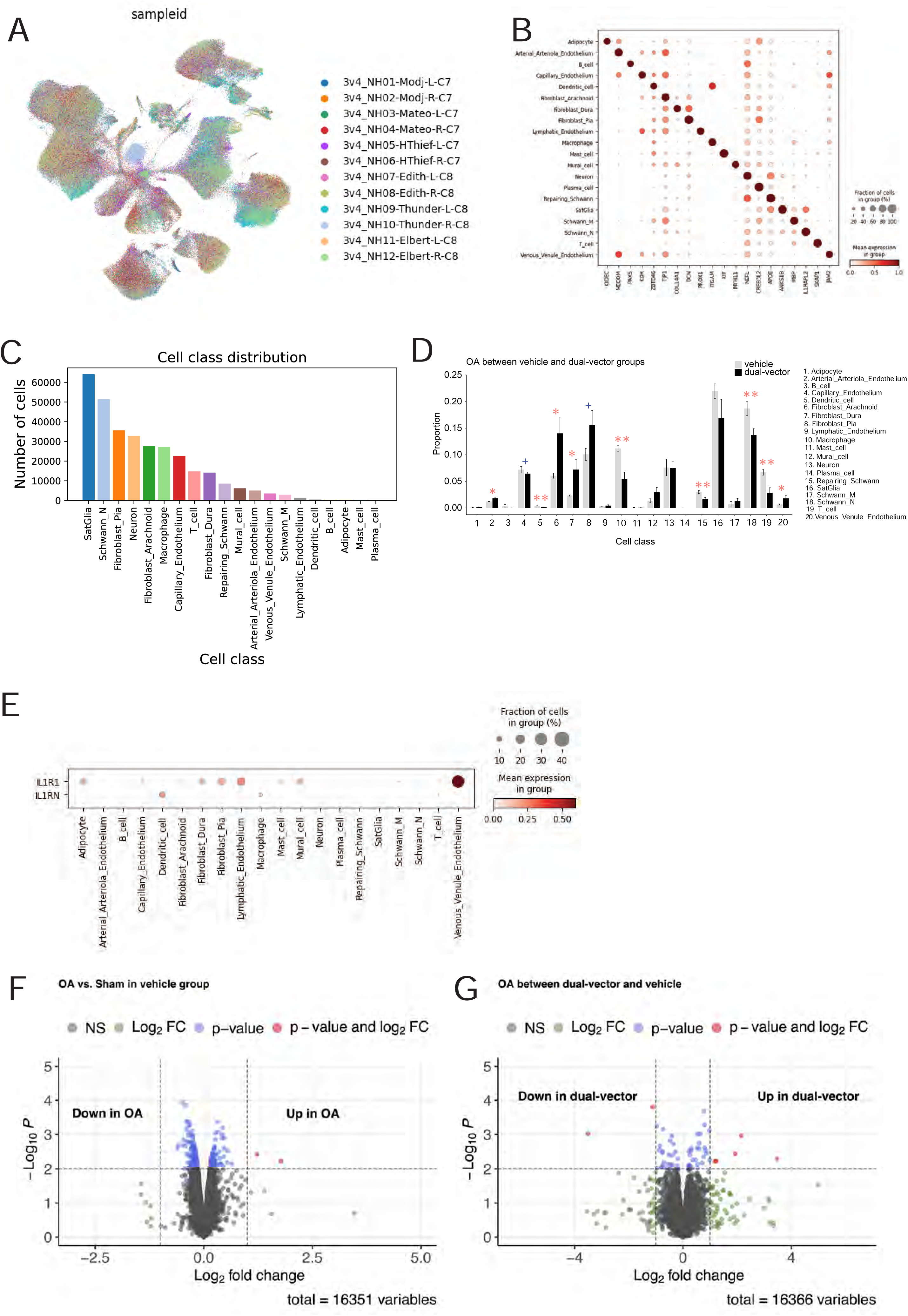
The cell atlas of the equine DRG. (A) UMAP showing the transcriptomic cell atlas derived from snRNA-seq data from 12 horse DRGs, with cells colored by sample ID. (B) Dot plot showing expression of marker genes across major cell classes in the snRNA-seq data. (C) Bar plot summarizing the cell abundance of each major cell class. (D) Bar plot showing the comparison of major class cell proportions in OA DRGs between the placebo and dual-vector groups. (E) Dot plot showing expression of *IL-1R1* and *IL-1RN* across major cell classes. (F) Volcano plot showing differential gene expression between OA and Sham DRGs in the placebo group based on bulk RNA-seq analysis. (G) Volcano plot showing differential gene expression between dual-vector treated and vehicle treated DRGs based on bulk RNA-seq analysis. A one-tailed Wilcoxon ranking sum test was conducted to calculate the P-values. +: P≤0.1, *: P≤0.05, **: P≤0.01, and ***: P≤0.001. Bar graphs depict mean values ± standard errors.

**Fig. S10.**
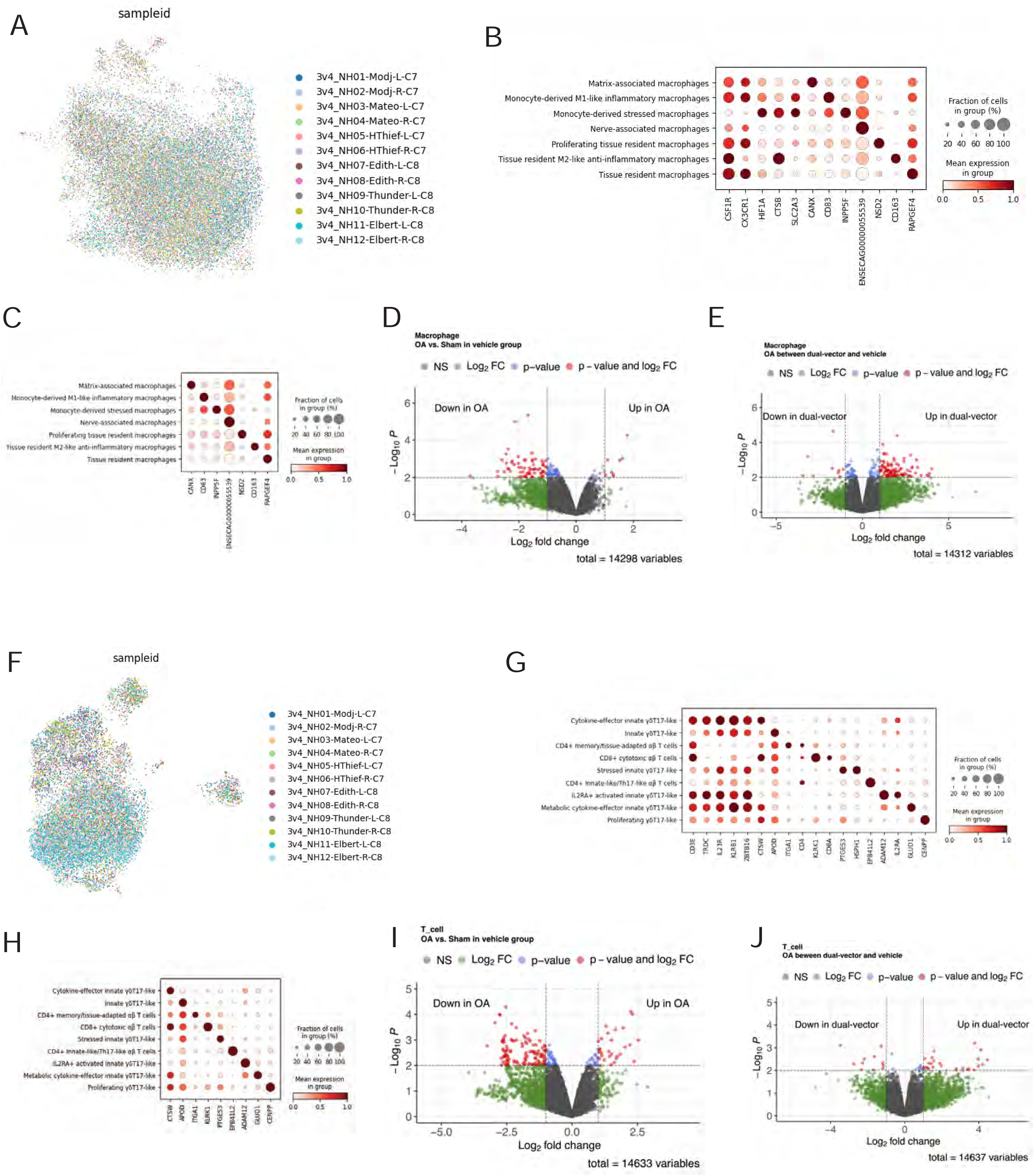
Macrophage and T cells populations in the equine DRG. (A) UMAP showing macrophages derived from snRNA-seq data from 12 horse DRGs, with cells colored by sample ID. (B) Dot plot showing expression of canonical marker genes across macrophage populations. (C) Dot plot showing expression of cluster-specific marker genes across macrophage populations. (D) Volcano plot showing differential gene expression between OA and Sham macrophages in the placebo group based on pseudo-bulk analysis. (E) Volcano plot showing differential gene expression between dual-vector treated and vehicle treated OA macrophages. (F) UMAP showing T-cells derived from snRNA-seq data from 12 horse DRGs, with cells colored by sample ID. (G) Dot plot showing expression of canonical marker genes across T-cell populations. (H) Dot plot showing expression of cluster-specific marker genes across T-cell populations. (I) Volcano plot showing differential gene expression between OA and sham T-cells in the placebo group. (J) Volcano plot showing differential gene expression between dual-vector treated and vehicle treated OA T cells based on pseudo-bulk analysis.

**Fig. S11.**
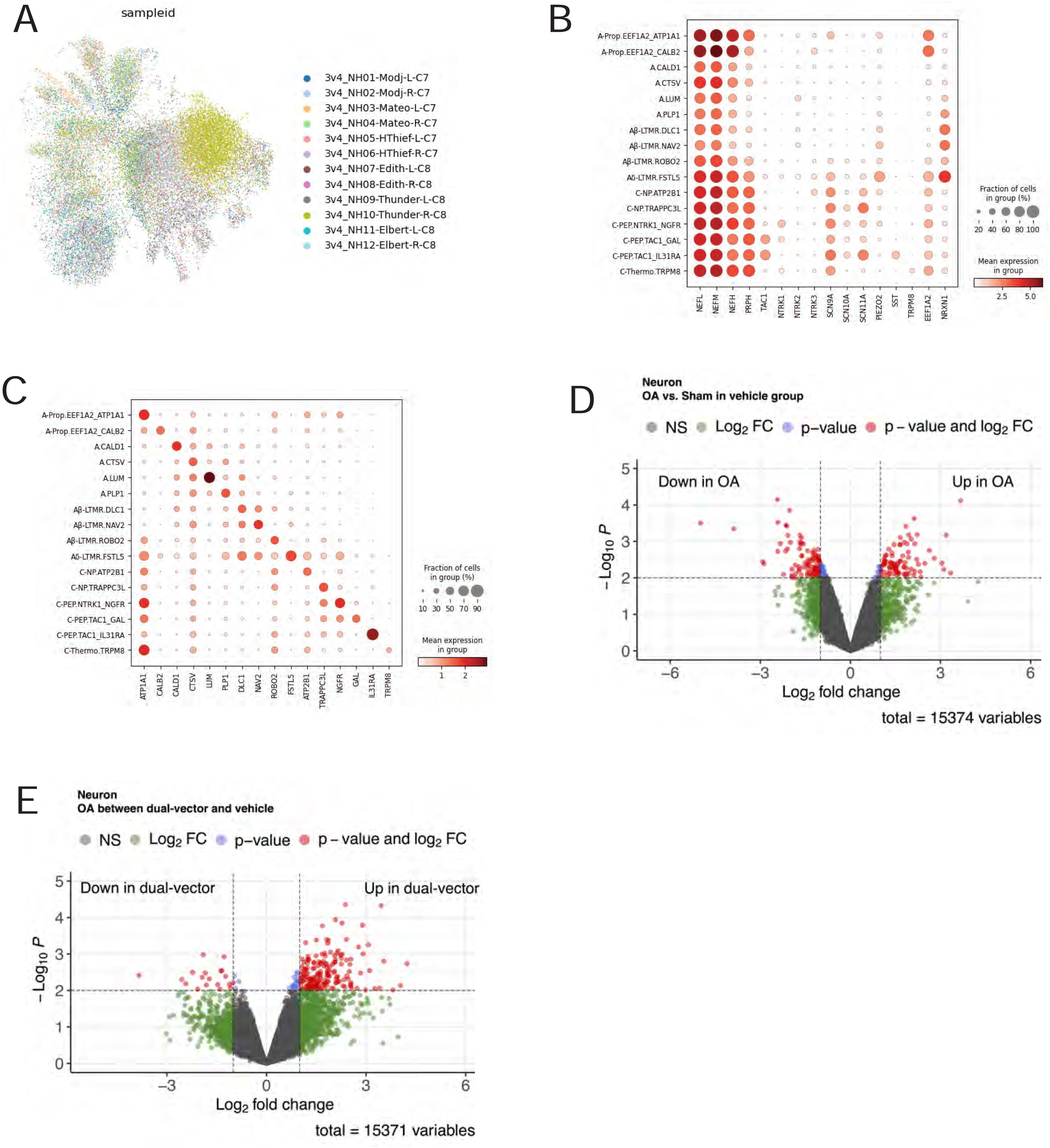
Neuronal populations in equine DRG. (A) UMAP showing neurons derived from snRNA-seq data from 12 horse DRGs, with cells colored by sample ID. (B) Dot plot showing expression of canonical marker genes across neuronal populations. (C) Dot plot showing expression of cluster-specific marker genes across neuronal populations. (D) Volcano plot showing differential gene expression between OA and sham neurons in the placebo group based on pseudo-bulk expression. (E) Volcano plot showing differential gene expression between dual-vector treated and vehicle treated OA neurons.

